# AAV-NRF2 protects retinal and choroidal vasculature in a GDF15-dependent manner in an oxidative damage model of AMD

**DOI:** 10.64898/2026.05.13.724735

**Authors:** Shuai Wang, Sophia Zhao, Adam Daniels, Efrat Naaman, Apolonia Gardner, Tianxi Wang, Ye Sun, Zhongjie Fu, Lois E. H. Smith, Constance L. Cepko

**Author notes:** Correspondence should be addressed to C.L.C.

## Abstract

Oxidative stress is proposed to be a driver of age-related diseases. Age-related macular degeneration is one such disease, where the retinal pigment epithelium (RPE) is affected early in the disease. Vasculature damage also occurs, sometimes preceding RPE damage. To model some aspects of dry AMD, we used the NaIO3 mouse model of oxidative damage. Disruption of the deep retinal vascular plexus, disorganization and death of capillaries within the choriocapillaris, and marked electroretinographic decline were observed. AAV overexpressing the transcription factor, NRF2, which induces anti-oxidation enzymes and represses inflammation, was tested for protection of damage. The BEST1 promoter limited expression to the RPE. The RPE, photoreceptors, and vascular architecture in both retinal and choroidal compartments were protected. Conditioned medium from RPE-choroid explants, infected by AAV8/BEST1-NRF2, was sufficient to transfer partial protection in vivo, indicating that NRF2 induces a protective secreted factor(s). Analysis of RNA-seq data identified growth differentiation factor 15 (GDF15) as a candidate downstream mediator. Injection of recombinant GDF15 reproduced key protective phenotypes in vivo, whereas Gdf15-deficiency attenuated NRF2-mediated rescue. Pharmacologic inhibition of TGF-β receptor signaling diminished NRF2 associated protection, supporting involvement of this signaling pathway. In a laser-induced choroidal neovascularization model, intravitreal GDF15 injection reduced fluorescein leakage and lesion size. These findings support a model in which NRF2 activation in the RPE induces expression of GDF15, which is capable of protecting the RPE, photoreceptors, and the retinal and choroidal vasculature. NRF2 and GDF15 have therapeutic potential for ocular diseases, as well as for other diseases with vascular pathology.

## INTRODUCTION

Age-related macular degeneration (AMD) is a leading cause of central vision loss in older adults. It is driven by a complex interplay of aging, genetic susceptibility, and environmental stressors (1, 2). Among the pathogenic mechanisms implicated in non-neovascular (dry) AMD, oxidative stress has been suggested to be a major driver. It is thought to first affect the retinal pigment epithelium (RPE) (3). Because the RPE supports photoreceptor metabolism, retinoid recycling, and barrier function, oxidative injury to this monolayer is considered a pivotal event in outer retinal failure (3-5). The RPE forms the outer blood-retinal barrier and lies at the interface between the retina and the vasculature that supplies the outer retina, the choroid. It regulates immune privilege, mediates the transport of nutrients and waste, provides the critical chromophore for vision, and provides trophic support (4, 5). Classic studies established that the RPE secretes VEGF in a polarized manner from its basal surface toward the choroid (6). The RPE-derived VEGF is required for choroidal vascular development and maintenance, underscoring the importance of the RPE for this adjacent vascular bed (7). Histopathologic studies of human AMD have further shown a close relationship between RPE abnormalities and choriocapillaris loss (8-12). Indeed, choriocapillaris dropout has been linked to drusen, accumulations of oxidized lipids and proteins that are a sign of AMD. In some analyses, choriocapillaris loss has been noted to precede other symptoms of AMD, supporting the view that vascular instability may be an early driver of AMD (9-11).

To better understand the relationship of the RPE, the choroid, and inner retinal vessels, it would be useful to have an animal model to study in health and disease. Systemic injection of sodium iodate (NaIO3) has been widely used to model oxidative stress. Its primary toxicity is to the RPE and it can reproduce key features of outer retinal degeneration relevant to dry AMD (13). Although this model has traditionally been used to study RPE loss and secondary photoreceptor damage, accumulating evidence indicates that NaIO3 injury also perturbs the choriocapillaris and outer retinal tissue environment (13). This makes it a useful framework for asking whether processes initiated in the RPE can influence vascular stability in a non-cell-autonomous manner.

NRF2 is a master transcriptional regulator that positively regulates antioxidant and cytoprotective gene programs (14, 15). In the eye, impaired NRF2 signaling has been linked to increased vulnerability of the aging RPE, and genetic disruption of Nrf2 in mice leads to several phenotypes of AMD (16-18). Conversely, our previous work has shown that AAV-mediated overexpression of NRF2 in the RPE using the BEST1 promoter preserves RPE structure in a mouse model of an inherited retinal degeneration (19). In the acute NaIO3 model in mice and in rats, the same AAV8/BEST1-NRF2 prevented damage to the RPE and photoreceptors, and preserved vision (20). Whether NRF2 overexpression in the RPE can also stabilize neighboring retinal and choroidal vasculature has not been addressed.

One reason to suspect a protective role of NRF2 on the vasculature is that it not only regulates detoxification and redox homeostasis, but also inflammatory and secretory programs (14, 15, 21). Among stress-induced secret factors, growth differentiation factor 15 (GDF15) is of particular interest. GDF15 is a divergent TGF- β superfamily cytokine induced by mitochondrial stress, inflammation, and tissue injury, and has been increasingly recognized as a context dependent mediator of adaptation and tissue protection (22, 23). It is directly regulated by NRF2, among other transcriptional regulators. Although the canonical metabolic effects of circulating GDF15 are mediated through the hindbrain receptor, GFRAL (24, 25), it has well documented actions in peripheral tissues (22, 23). The mechanism of its action and the receptor(s) it uses more peripherally are still unclear. In ocular systems, GDF15 is induced by retinal injury, can protect retinal ganglion cells, and has recently been implicated in limiting autoimmune uveitis through modulation of retinal microglial responses (26-28). These observations make GDF15 a plausible candidate for stress responsive signaling within the injured retina, but its potential role downstream of RPE-specific NRF2 overexpression, particularly in regulating ocular vascular stability, has not been examined. Here, we found that RPE-specific overexpression using AAV led to protection of choroidal and retinal vessels. The protection was partially dependent upon GDF15. Injection of recombinant GDF15 was found to recapitulate key aspects of NRF2-mediated protection, including preservation of retinal and choroidal vasculature, as well as RPE and photoreceptor structure.

## RESULTS

### NaIO3 Induces Progressive Retinal and Choroidal Vascular Abnormalities with Functional Decline

To define vascular and tissue changes induced by oxidative injury, adult mice were systemically administered NaIO3 and analyzed longitudinally by fundus fluorescein angiography (FFA), electroretinography (ERG), and flat-mount imaging of retinal and RPE-choroid tissues (Fig 1A). Consistent with previous reports of NaIO3 injection (29), FFA revealed progressively increased hyperfluorescent signals over time, becoming more pronounced from 1 day to 1 week, and further increasing by 4 weeks after injury. Quantification of mean fluorescein signal intensity confirmed a significant treatment and time-dependent increase in the angiographic abnormalities in NaIO3-treated eyes (Fig. 1B, F). Functional testing further showed that NaIO3 injury was associated with substantial retinal dysfunction. ERG measurements demonstrated reduced scotopic and photopic responses, as well as impaired oscillatory potentials, at 4 weeks in NaIO3-treated versus saline-treated controls (Fig. 1C, G).

**Figure 1.**
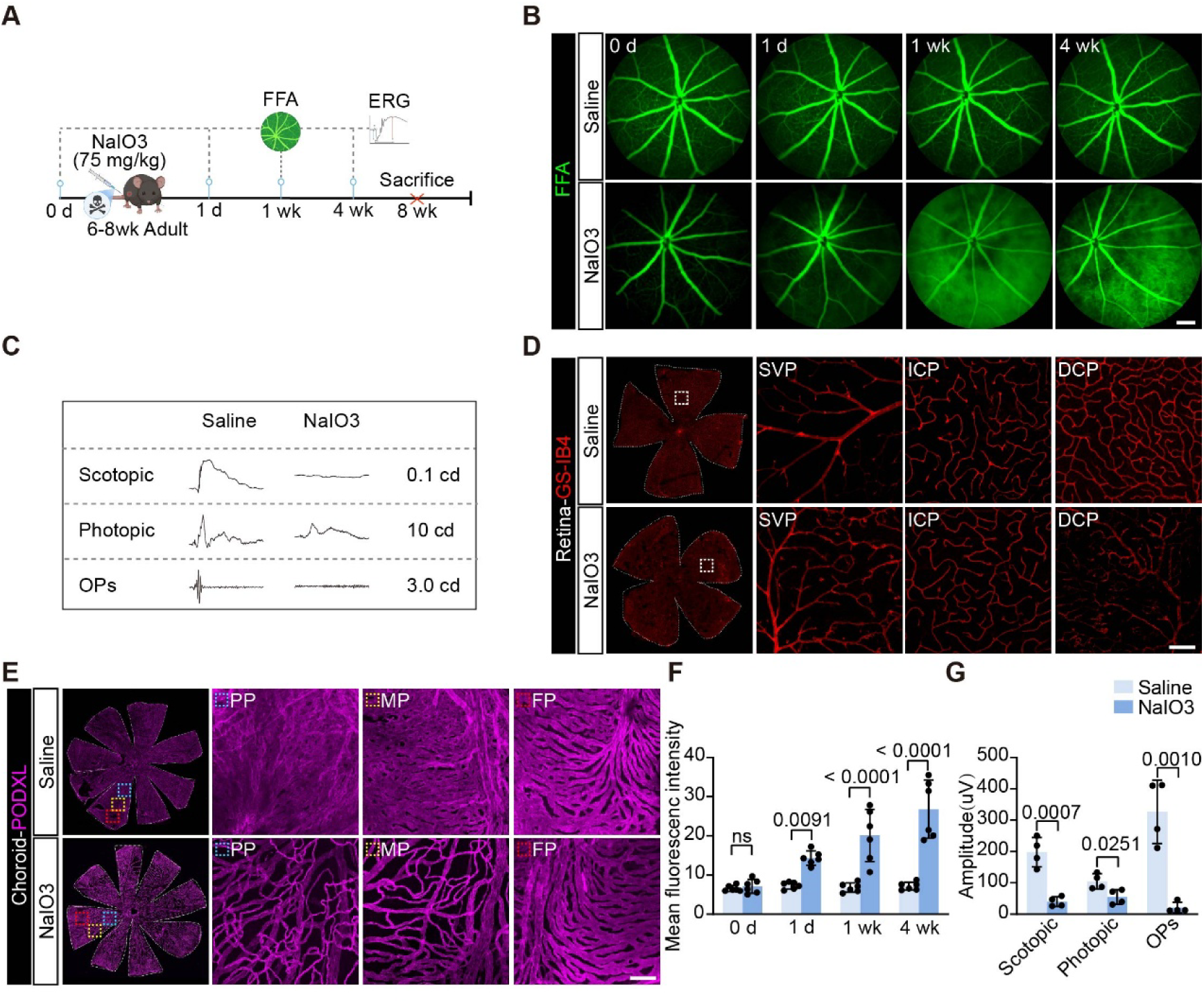
Characterization of retinal and choroidal changes following NaIO3 administration. **(A)** Experimental timeline illustrating systemic NaIO3 administration in adult mice. **(B)** Fundus fluorescein angiography (FFA) images at the indicated time points. Scale bar, 100 μm. **(C)** Representative electroretinography (ERG) recordings showing scotopic and photopic, oscillatory potentials (OPs) at 4 weeks. **(D)** GS-IB4-stained retinal flat-mount. Insets show higher-magnification views of the superficial vascular plexus (SVP), intermediate capillary plexus (ICP), and deep capillary plexus (DCP). Scale bar, 100 μm. **(E)** Representative podocalyxin-stained RPE–choroid flat-mount images illustrating choroidal vascular organization. Insets show higher-magnification views of the posterior pole (PP, blue), mid-periphery (MP, yellow), and far periphery (FP, red). Scale bar, 100 μm. **(F)** Quantification of mean fluorescein intensity shown in B (n =6 eyes per group, mean±SD, two-way ANOVA with Šídák’s multiple-comparisons test). **(G)** Quantification of ERG waves shown in C (n =4 eyes per group, mean ± SD, unpaired t-test).

Retinal vessels were visualized in flat mounts using GS-IB4, a lectin commonly used to label endothelial cells and vascular networks. The superficial vascular plexus (SVP), intermediate capillary plexus (ICP), and deep capillary plexus (DCP) were all well organized in control retinas. Following NaIO3 treatment, the SVP and ICP remained largely preserved, with no obvious alterations in overall vascular organization. In contrast, the DCP exhibited reproducible structural abnormalities, including irregular vascular trajectories and narrowed luminal profiles, indicating selective vulnerability of the DCP network (Fig. 1D). Because the RPE is immediately adjacent to the choriocapillaris, the choroidal vasculature was examined. In control eyes, the choroidal vasculature exhibited distinct regional patterns: in the central posterior pole (PP), the choriocapillaris formed a dense honeycomb-like meshwork; in the mid-peripheral (MP) region, the capillary network displayed a more linear arrangement with fewer interconnections and larger intercapillary spaces; and in the far periphery (FP), choroidal vessels showed a characteristic palm-shaped pattern. Following NaIO3 treatment, these organized vascular patterns became disrupted (Fig. 1E). In the PP, the normally dense choriocapillaris meshwork became fragmented with reduced capillary density. In the MP region, the vascular network lost its compact organization and appeared sparser and more atrophic, with enlarged intervascular spaces, while in the FP there was little disruption. Together, these data define the vascular and functional phenotypes of the NaIO3 injury model. These include progressive angiographic abnormalities, retinal functional decline, DCP changes, and prominent choroidal vascular disruption, consistent with prior reports of NaIO3-induced angiographic changes and choriocapillaris atrophy (29-31).

### RPE-Specific NRF2 Overexpression Preserves Ocular Vasculature

Given our previous study showing that NRF2 overexpression prevented damage to both RPE and photoreceptor cells following NaIO3 injection (20), it was of interest to determine if it could also preserve ocular vascular architecture. To achieve RPE-specific expression, an AAV with the RPE-specific BEST1 promoter was used to drive NRF2 expression (19). As a control, an AAV using the same promoter but lacking a functional open reading frame (AAV8/BEST1-6xSTOPmutGFP, hereafter AAV-CTL) was used. These were delivered by subretinal injection at postnatal day 0 (P0), together with a small amount of vector (AAV8/RedO-H2B-GFP, hereafter AAV-GFP) to trace the infected area (20). Mice were allowed to mature, and vascular analyses were performed in adulthood following NaIO3 administration. FFA revealed that eyes expressing AAV8/BEST1-NRF2 exhibited reduced angiographic abnormalities compared with AAV-CTL injected eyes after NaIO3 injection (Fig. 2B, E). To further assess whether NRF2 could also protect the mature eye, AAV8/BEST1-NRF2 was injected subretinally into adult mice. NaIO3 was injected intraperitoneally, and FFA was performed. Because adult subretinal injection typically yields partial transduction, this paradigm provides localized, rather than global transduction (20). AAV8/BEST1-NRF2 reduced the hyperfluorescent signal in a partial and spatially restricted manner, with the effect most evident in GFP-positive regions (Fig. S1). These data indicate that even when delivered to fully developed eyes, NRF2 is sufficient to confer localized vascular protection.

**Figure 2.**
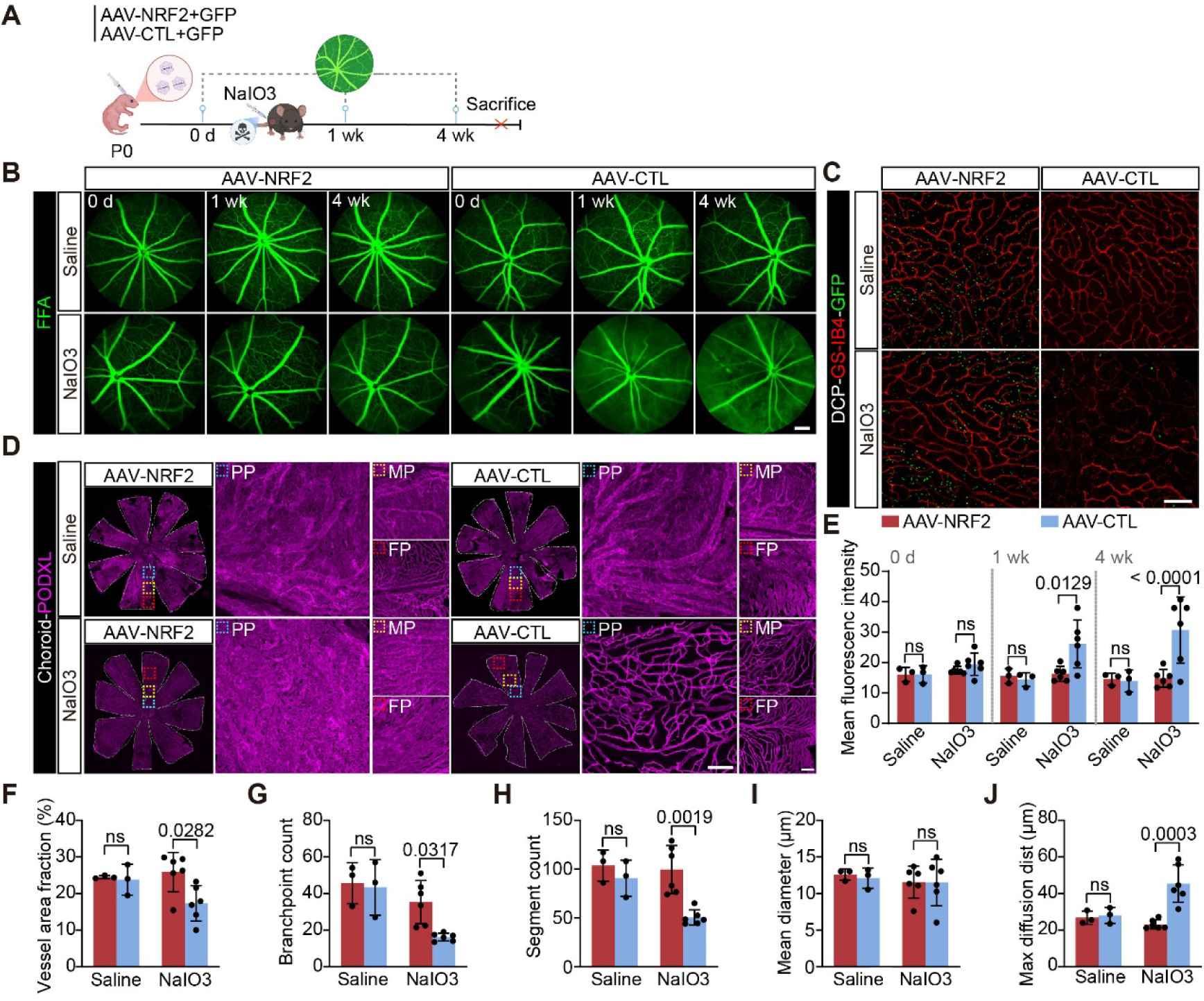
NRF2 expression in RPE preserves retinal and choroidal vasculature. **(A)** Experimental design illustrating P0 subretinal delivery of AAV8/BEST1-NRF2 or AAV-CTL together with a GFP tracer vector, followed by saline or NaIO3 administration in adulthood. **(B)** FFA images from AAV8/BEST1-NRF2 and AAV-CTL eyes following saline or NaIO3 administration. Scale bar, 100 μm. **(C)** GS-IB4-stained retinal flat-mount images showing the DCP. GFP indicates AAV-transduced retina. Scale bar, 100 μm. **(D)** Representative podocalyxin stained RPE–choroid flat-mount images. Higher magnification views of choroidal vasculature in PP (blue), MP (yellow), and FP (red). Scale bar, 100 μm. **(E)** Quantification of mean fluorescein intensity from FFA images shown in B (n = 3 eyes per group for saline and n = 6 eyes per group for NaIO3, mean ± SD, two-way ANOVA with Šídák’s multiple-comparisons test). **(F-J)** Quantification of retinal vascular parameters in the DCP shown in C (n = 3 eyes per group for saline and n = 6 eyes per group for NaIO3, mean ± SD, two-way ANOVA with Šídák’s multiple-comparisons test).

To further evaluate retinal microvascular preservation, the DCP was examined in GS-IB4-stained retinal flat mounts. In AAV-CTL infected eyes, NaIO3 treatment caused disruption and simplification of DCP architecture. Eyes infected with AAV8/BEST1-NRF2 retained a denser and more continuous capillary network (Fig. 2C). GFP signal was detected broadly across the retina in these neonatal-injected samples, consistent with the extensive infection that is typical of neonatal injections. Quantitative analysis demonstrated that vascular area fraction, branchpoint, and segment number were higher in the AAV8/BEST1-NRF2 group, whereas maximal diffusion distance was lower compared with NaIO3-treated AAV-CTL infected eyes. A lower maximal diffusion distance indicates fewer avascular spaces within the analyzed DCP field, consistent with improved vascular coverage and oxygen diffusion capacity. We next examined whether AAV8/BEST1-NRF2 infection also preserved the choroidal vasculature. In AAV-CTL infected eyes in animals injected with NaIO3, the PP region showed marked loss and fragmentation of the normally dense choriocapillaris meshwork. In contrast, AAV8/BEST1-NRF2 eyes retained a more continuous vascular structure in this region. A similar pattern was observed in the MP region, where AAV-CTL infected eyes displayed a looser vascular network with enlarged intercapillary spaces, whereas AAV8/BEST1-NRF2 infected eyes maintained a denser vascular organization (Fig. 2D). Together, these findings indicate that RPE targeted NRF2 overexpression preserves both retinal and choroidal vascular architecture following NaIO3-induced injury. The adult injection FFA data further indicate that NRF2 can provide vascular protection in mature eyes.

### AAV8/BEST1-NRF2 Induced Secreted factor (s) Prevents Vascular Damage

To determine whether there was a secreted factor(s) produced by the RPE infected with AAV8/BEST1-NRF2 which confers vascular protection, conditioned medium(CM) was generated by RPE-choroid explants derived from mice infected with AAV8/BEST1-NRF2 or the AAV-CTL vector (NRF2-CM and CTL-CM, respectively) (Fig. 3A). Explants were made from eyes infected in vivo at P0, and then aged to 8 weeks. The freshly explanted tissue was cultured for 24 h, and the collected supernatants were fractionated into >10 kDa and <10 kDa components using centrifugal filtration. Each fraction was delivered by subretinal injection into adult mice before NaIO3 administration, together with a GFP tracer virus. FFA performed 1 day after NaIO3 administration revealed that both the >10 kDa and <10 kDa fractions of NRF2-CM reduced hyperfluorescent signal compared with the corresponding CTL-CM fractions (Fig. 3B). The protective effect was localized to GFP-positive regions. Because both size fractions displayed protective activity, it was of interest to determine which size fraction had a higher concentration of protective activity. To this end, both size fractions of the NRF2-CM were diluted tenfold before subretinal delivery. The >10KDa diluted material retained protective activity, whereas the <10 kDa fraction lost detectable protection, consistent with the idea that the protective activity is a protein (Fig. S2A).

**Figure 3.**
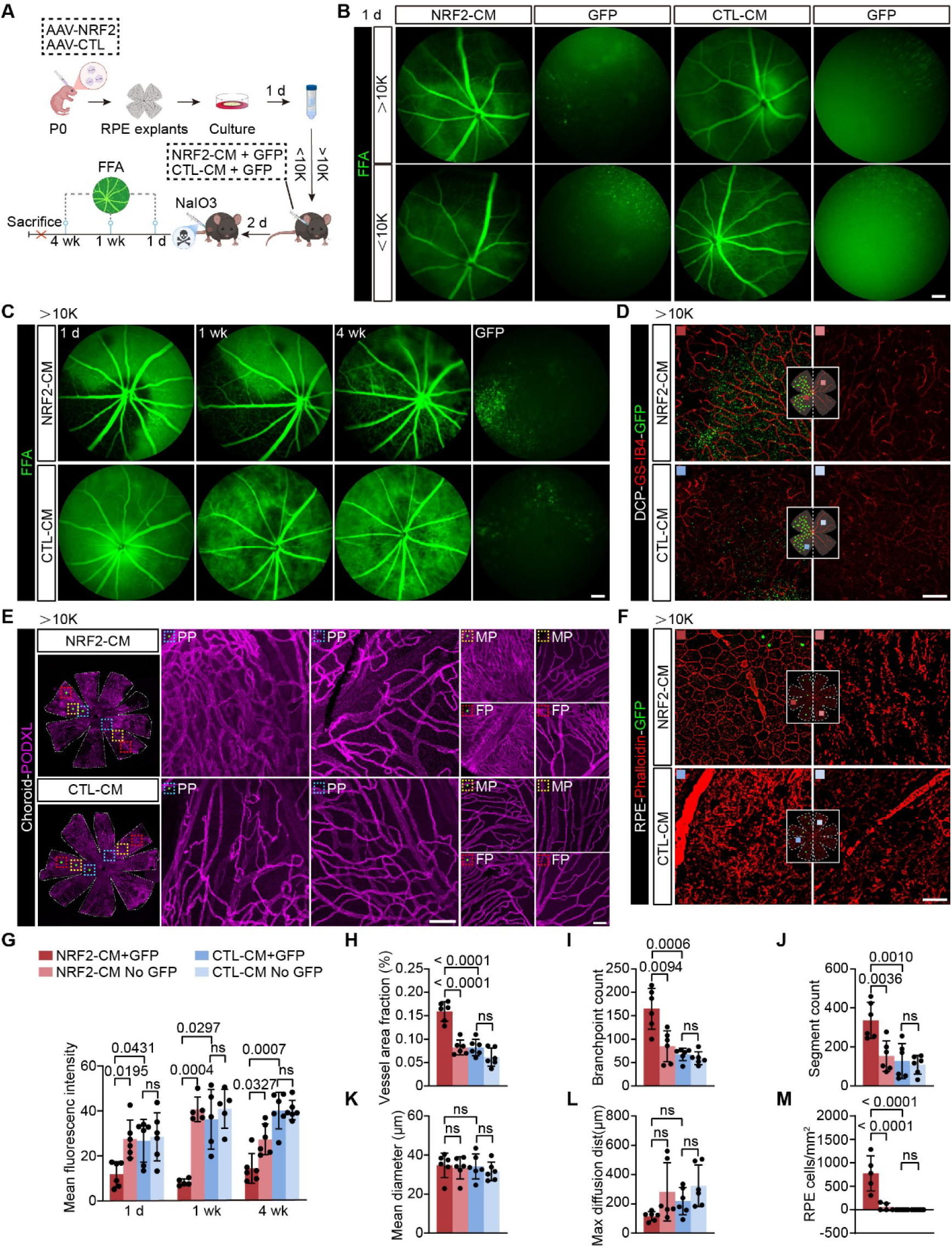
NRF2 Conditioned Medium transfers partial vascular protection in vivo. **(A)** Schematic illustration of conditioned medium (CM) preparation from RPE–choroid explants and subsequent in vivo transfer experiment. **(B)** FFA images at 1 day after NaIO3 administration following subretinal injection of NRF2-CM or CTL-CM fractions (>10 kDa or <10 kDa) together with GFP tracer. **(C)** Longitudinal FFA images following subretinal delivery of the >10 kDa NRF2-CM or CTL-CM together with GFP tracer. Scale bar, 100 μm. **(D)** Retinal flat-mounts showing the DCP. GFP signal marks the injected retinal region. Scale bar, 100 μm. **(E)** Representative podocalyxin-stained RPE–choroid flat-mounts. Higher magnification views of choroidal vasculature in PP (blue), MP (yellow), and FP (red). Scale bar, 100 μm. **(F)** Phalloidin-stained RPE flat-mounts showing RPE morphology. Scale bar, 50 μm. **(G)** Quantification of FFA shown in C (n = 6 eyes per group, mean ± SD, two-way ANOVA with Tukey’s multiple-comparisons test). **(H-L)** Quantitative analysis of retinal vascular parameters shown in D (n = 5 eyes per group, mean ± SD, one-way ANOVA with Tukey’s multiple-comparisons correction). (**M)** Quantification of RPE cell counts shown in F (n = 5 eyes per group for NRF2 and n = 7 eyes for CTL, mean ± SD, one-way ANOVA with Tukey’s multiple-comparisons correction).

Subsequent analyses were therefore conducted on the >10 kDa fraction. Longitudinal FFA showed that eyes receiving NRF2-CM exhibited markedly reduced diffuse hyperfluorescence at 1 day, 1 week, and 4 weeks after injury compared with eyes injected with CTL-CM (Fig. 3C). Quantification confirmed a significant reduction in fluorescein signal intensity in NRF2-CM-treated eyes (Fig. 3G). To directly examine retinal vascular architecture, retinal flat mounts were stained with GS-IB4. In GFP-positive regions, NRF2-CM-treated eyes displayed a more continuous and organized DCP than CTL-CM-treated eyes, whereas GFP-negative regions showed comparable vascular disruption between groups (Fig. 3D). Quantitative analysis of the DCP revealed increased vascular density, branchpoint number, and total vessel length, together with a reduction in maximal diffusion distance in NRF2-CM-treated retinas relative to CTL-CM controls. These observations demonstrated that NRF2-CM protects retinal vascular integrity (Fig. 3H-L). The choroidal vasculature also was examined. In GFP-positive regions, NRF2-CM-treated eyes preserved a denser and more organized choriocapillaris network than CTL-CM treated eyes, although this preservation was less complete than that observed following infection with AAV8/BEST1-NRF2. In contrast, GFP-negative regions displayed similar vascular disruption in both groups (Fig. 3E). RPE morphology was assessed using phalloidin staining. Within GFP-positive regions, NRF2-CM-treated eyes showed partial preservation of the RPE cellular mosaic compared with CTL-CM-treated eyes, although the morphology was slightly less regular than that observed in uninjured tissue (Fig. 3F). Together, these findings indicate that the RPE infected by AAV8/BEST1-NRF2 secretes at least one factor that is sufficient to confer partial, spatially restricted vascular protection against oxidative injury.

### Identification of Candidate Secreted Factors Downstream of NRF2

To identify candidate mediators of the vascular protective activity of the NRF2-CM, RNA-seq data generated from RPE infected by AAV8/BEST1-NRF2 (19) was analyzed for upregulated secreted factors. This dataset was collected from infection of CD1 wild-type animals that were not treated with NaIO3 (Fig. 4A). To prioritize candidates capable of signaling to vascular cells, upregulated genes were compared with a list of receptors that are in non-neuronal retinal cells, including vascular endothelial cells (32). By cross-referencing NRF2 induced secreted ligands with receptor expression across retinal and choroidal endothelial populations, we identified GDF15 and oncostatin M (OSM) as candidate mediators whose predicted receptors are expressed in vascular endothelial cells (Fig. 4B).

**Figure 4.**
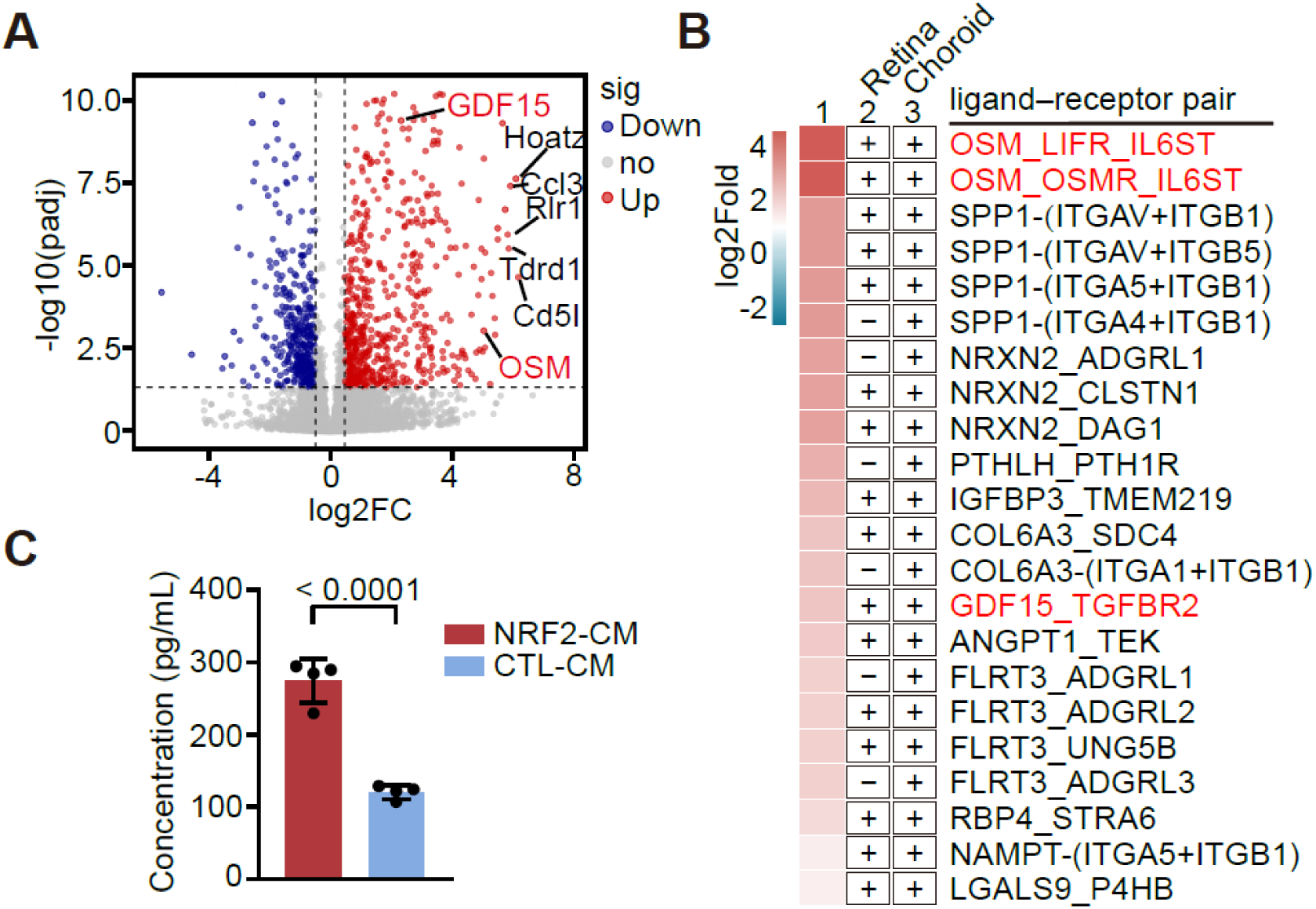
Identification of GDF15 as a candidate secreted effector induced by RPE NRF2 overexpression. **(A)** Analysis of the RNA-seq dataset (19). Identified secreted factors upregulated in the RPE–choroid complex following AAV8/BEST1-NRF2 relative to AAV-BEST1-GFP infection. Analysis was on RPE samples from CD1 wild-type mice. **(B)** Ligand–receptor prioritization of NRF2-induced secreted ligands. Upregulated candidate secreted ligands from the RPE bulk RNA-seq dataset were cross-referenced with receptor expression in retinal and choroidal endothelial cells reported in a non-neuronal retinal single-cell RNA-seq atlas (32). Each row shows a candidate ligand–receptor pair for which the corresponding receptor was detected in retinal endothelial cells, choroidal endothelial cells, or both. The heatmap in column 1 indicates the ligand log2 fold change in NRF2-expressing RPE cells from bulk RNA-seq data (19). Columns 2 and 3 indicate whether the corresponding receptor was detected in retinal or choroidal endothelial cells, respectively, based on single-cell RNA-seq data (32). Red text highlights the two candidate ligands selected for functional testing, OSM and GDF15. **(C)** ELISA quantification of GDF15 protein levels in unconcentrated NRF2-CM or CTL-CM (n =4 eyes per group, mean ± SD, t-test).

We tested whether either factor could reproduce the vascular protective phenotype in vivo. Recombinant OSM or GDF15 was delivered subretinally before NaIO3 injury, and FFA analyses at 1 day and 1 week post NaIO3 injury were performed. OSM showed no significant protection compared with PBS treated controls (Fig. S2B and C). However, GDF15 showed protection. An ELISA was carried out to determine if GDF15 was present in the CM, and if it was at a higher level in NRF2-CM relative to CTL-CM. The ELISA showed that indeed, this was the case (Fig. 4C). Together, these findings identify GDF15 as induced by AAV8/BEST1-NRF2 and nominate it as a candidate mediator of NRF2-dependent protection.

### Recombinant GDF15 Partially Recapitulates AAV8/BEST1-NRF2-induced Vascular and Tissue Protection

To test whether GDF15 is sufficient to produce features of AAV8/BEST1-NRF2-mediated protection, adult mice received subretinal injections of recombinant mouse GDF15 (rGDF15, 10 μM, 1 μL/eye) (28) or PBS, together with an AAV-GFP tracer, followed by NaIO3 administration 2 days later (Fig. 5A). Serial FFA showed that rGDF15 treated eyes exhibited reduced hyperfluorescent abnormalities compared with PBS treated eyes at 1 day, 1 week, and 4 weeks after injury (Fig. 5B). Comparison of GFP-positive and GFP-negative regions showed that the protective effect was localized to the injected area, indicating a local action of rGDF15 (Fig. 5B). To further assess retinal vascular structure, retinal flat mounts stained with GS-IB4 were analyzed. In the DCP, rGDF15 treated GFP-positive regions retained a more continuous capillary network, whereas PBS treated eyes displayed pronounced capillary disruption and vessel rarefaction following NaIO3 injury (Fig. 5C). Quantitative analyses demonstrated increased vessel area fraction, vessel length, branchpoint number, and segment count in rGDF15-treated retinas compared with PBS controls, along with reduced maximal diffusion distance, indicating improved vascular integrity and oxygen delivery capacity (Fig. 5 H-J and Fig. S3).

**Figure 5.**
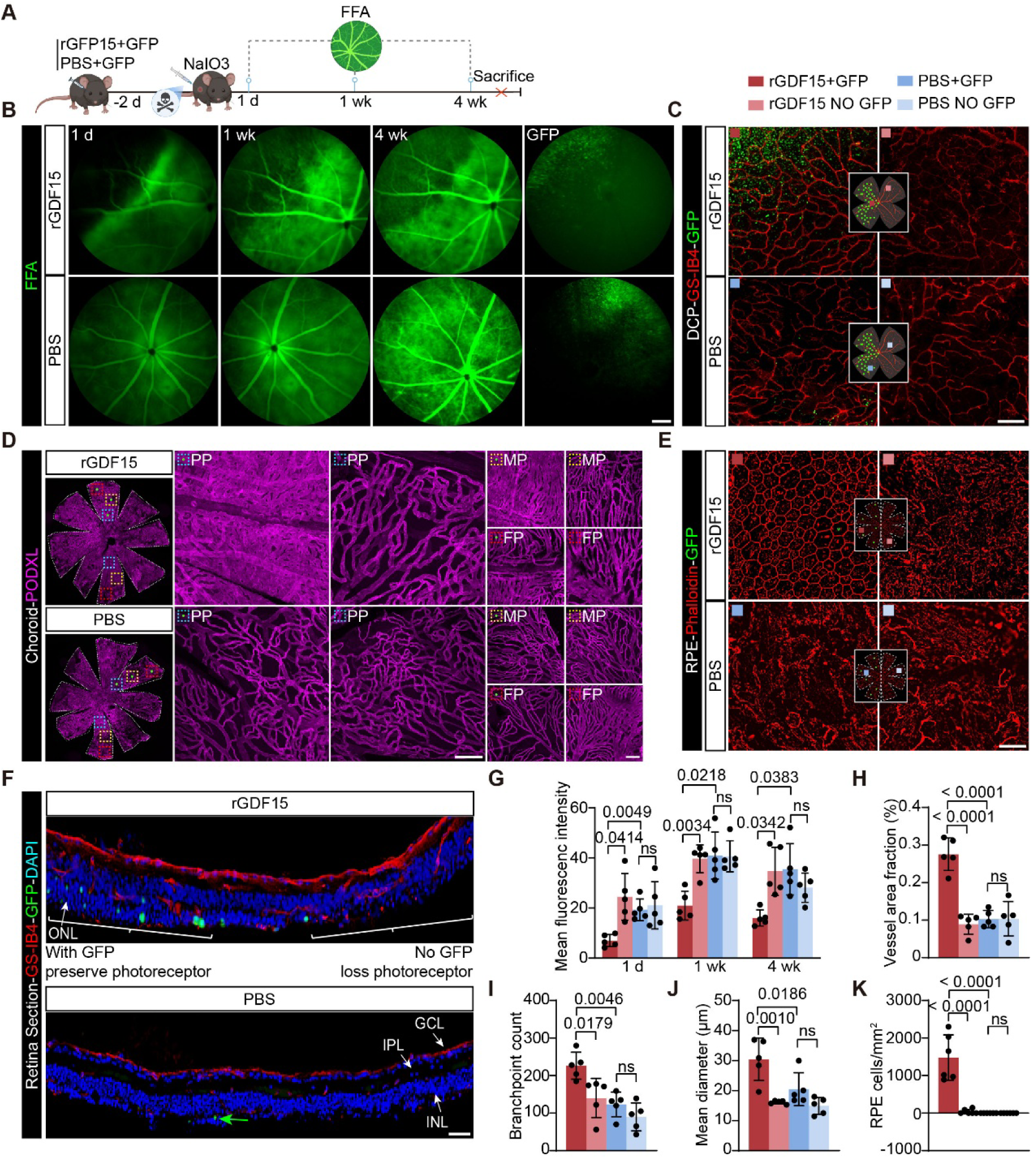
rGDF15 partially recapitulates NRF2-CM mediated vascular protection. **(A)** Experimental design. Adult mice received a subretinal injection of recombinant mouse GDF15 (rGDF15, 10 μM, 1 μL/eye) or PBS together with a GFP tracer. NaIO3 was administered 2 days later. **(B)** FFA images over time after NaIO3 administration. GFP images indicate the injected region. Scale bar, 100 μm. **(C)** GS-IB4-stained retinal flat-mounts showing DCP. Scale bar, 100 μm. **(D)** Representative podocalyxin stained choroidal vasculature in PP (blue), MP (yellow), and FP (red). Scale bar, 100 μm. **(E)** Phalloidin stained RPE flat-mounts showing RPE morphology in rGDF15 and PBS treated eyes. Scale bar, 50 μm. **(F)** Retinal sections showing photoreceptor structure in GFP-positive and GFP-negative regions. Scale bar, 50 μm. **(G)** Quantification of mean fluorescein intensity in GFP-positive and GFP-negative regions shown in B (n = 5 eyes per group, mean ± SD, two-way ANOVA with Tukey’s multiple-comparisons test). **(H–J)** Quantification of retinal vascular parameters shown in C (n = 5 eyes per group, mean ± SD, one-way ANOVA with Tukey’s multiple-comparisons correction). **(K)** Quantification of RPE cell counts shown in E (n = 6 eyes per group, mean ± SD, one-way ANOVA with Tukey’s multiple-comparisons correction).

The choroidal vasculature was examined using podocalyxin immunohistochemistry on RPE-choroid flat mounts. In PBS-treated eyes, NaIO3 injury caused substantial disruption of the choriocapillaris network, with reduced capillary density and loss of the characteristic vascular architecture across central and peripheral regions. In contrast, rGDF15 treated eyes preserved a denser and more organized choriocapillaris network, particularly in the PP and MP regions (Fig. 5D).

To determine if rGDF15 might also prevent RPE damage, RPE and photoreceptor morphologies were assessed by immunohistochemistry. In rGDF15 treated GFP-positive regions, the RPE monolayer retained a relatively intact hexagonal cellular mosaic. In contrast, PBS-treated retinas exhibited severe disruption of RPE organization, with irregular cell boundaries and loss of the typical epithelial structure. Adjacent GFP-negative regions in rGDF15-treated eyes similarly showed marked RPE degeneration following NaIO3 injury (Fig. 5E). Retinal sections revealed that photoreceptor structure and number were better preserved in rGDF15 injected GFP-positive regions. In PBS-treated eyes, NaIO3 injury resulted in substantial thinning of the outer nuclear layer reflecting loss of photoreceptor cells. In contrast, rGDF15 treated regions maintained a thicker photoreceptor layer, whereas GFP-negative regions showed photoreceptor degeneration similar to PBS controls (Fig. 5F).

To explore the effective dose range, mouse rGDF15 was delivered subretinally at 50, 10, or 1 ng per eye before NaIO3 injury. The 50 ng dose reduced FFA hyperfluorescence at 1 day after injury, whereas 10 and 1 ng did not show clear protection at this early time point (Fig. S4). By 1 week, however, protection became detectable in the lower-dose groups, indicating that rGDF15-mediated protection can occur at lower doses but with delayed onset.

Together, these results show that mouse rGDF15 reduces NaIO3-induced angiographic abnormalities of both the retinal and choroidal vasculature. In addition, rGDF15-treated regions showed better preservation of RPE cells and photoreceptors.

### Loss of GDF15 Partially Attenuates NRF2-Mediated Functional and Vascular Protection

The rGDF15 injections demonstrated that rGDF15 was sufficient to produce several of the protective features induced by AAV8/BEST1-NRF2 after NaIO3 injury. To determine whether GDF15 is necessary for the protective effects of AAV8/BEST1-NRF2, we examined Gdf15⁻/⁻ mice infected by AAV8/BEST1-NRF2 and subjected to NaIO3-injection (Fig. 6A). FFA showed that the reduction in hyperfluorescence observed in wild-type mice infected with AAV8/BEST1-NRF2-infected eyes was diminished in Gdf15⁻/⁻ mice. Quantification of FFA intensity confirmed attenuation of the NRF2 protective effect in the absence of GDF15 (Fig. 6B, C). To determine if loss of GDF15 might affect retinal physiology, the Gdf15⁻/⁻ mice, without AAV infection or NaIO3 injection, were tested by ERG. Baseline ERG responses were comparable between wild-type and Gdf15⁻/⁻ mice across scotopic, photopic, and oscillatory potential measurements (Fig. S5A). We therefore examined whether loss of GDF15 altered the functional benefit conferred by AAV8/BEST1-NRF2 after NaIO3 injury. ERG analysis showed that infection by AAV8/BEST1-NRF2 partially preserved retinal function in wild-type animals following NaIO3 treatment, as we previously reported (20) (Fig. S5B,6D). Under scotopic conditions, AAV8/BEST1-NRF2 infected wild-type eyes exhibited increased b-wave amplitudes compared with control eyes. In contrast, this improvement was significantly attenuated in Gdf15⁻/⁻ mice, in which AAV8/BEST1-NRF2 infection produced only a modest and non-significant increase in scotopic responses. Photopic b-wave responses were not significantly altered by AAV8/BEST1-NRF2 infection in either genotype, indicating limited effects on cone mediated responses under these conditions. In contrast, oscillatory potentials, which reflect inner retinal activity, showed the most pronounced AAV8/BEST1-NRF2 dependent improvement. AAV8/BEST1-NRF2 infection significantly preserved oscillatory potential amplitudes in wild-type mice after NaIO3 injury, whereas this effect was largely lost in Gdf15-deficient animals, consistent with a contribution of GDF15 to the functional benefit of NRF2, particularly in the inner retina.

**Figure 6.**
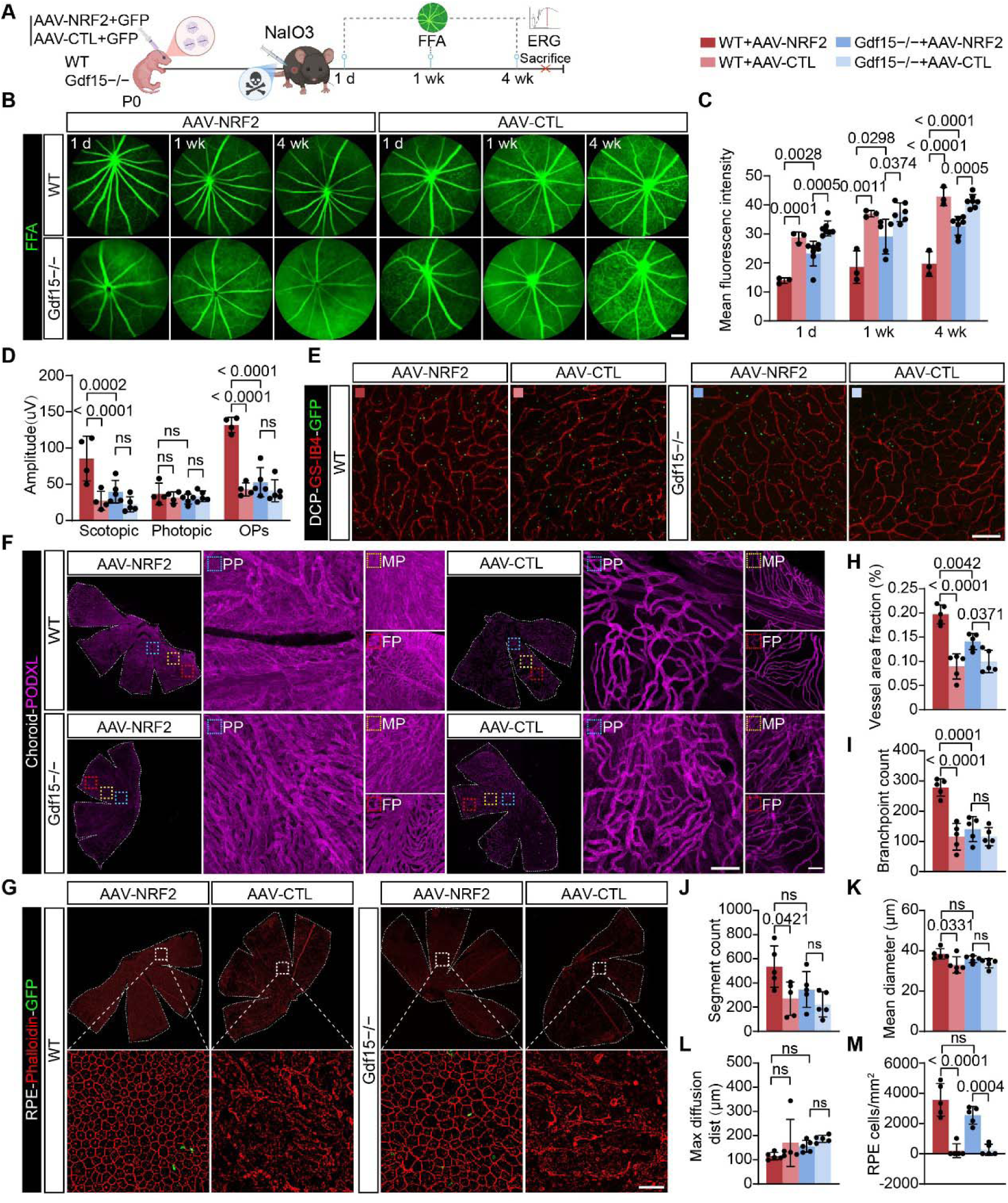
Loss of GDF15 attenuates NRF2-mediated protection following NaIO3-induced injury. **(A)** Experimental design. Wild-type and Gdf15⁻/⁻ mice received neonatal subretinal injection of AAV8/BEST1-NRF2 or AAV-CTL. In adulthood, mice were subjected to NaIO3-induced oxidative injury. Functional and structural analyses were performed at the indicated time points. **(B)** FFA images from the indicated groups following NaIO3 administration. Scale bar, 100 μm. **(C)** Quantification of fluorescein intensity shown in B (n = 3 eyes per group for Wild-type, n =7 eyes per group for Gdf15⁻/⁻ mice, mean ± SD, two-way ANOVA with Tukey’s multiple-comparisons test). **(D)** Quantification of ERG responses shown in Figure S5B (n = 4 eyes per group for wild-type, n = 5 eyes per group for Gdf15⁻/⁻, mean ± SD, two-way ANOVA with Tukey’s multiple-comparisons test). **(E)** GS-IB4 stained retinal flat-mounts showing DCP. Scale bar, 100 μm. **(F)** Representative podocalyxin-stained RPE–choroid flat-mounts showing choriocapillaris morphology in PP (blue), MP (yellow), and FP (red). Scale bar, 100 μm. **(G)** Phalloidin-stained RPE flat-mounts showing RPE. Scale bar, 50 μm. **(H–L)** Quantification of retinal vascular parameter shown in E (n = 5 eyes per group, mean ± SD, one-way ANOVA with Tukey’s multiple-comparisons correction). **(M)** Quantification of RPE cell counts shown in G (n = 5 eyes per group, mean ± SD, one-way ANOVA with Tukey’s multiple-comparisons correction).

To examine retinal vascular architecture, retinal flat mounts were analyzed. In wild-type retinas, AAV8/BEST1-NRF2 infection preserved a more continuous capillary network in the DCP after injury. In contrast, Gdf15-deficient retinas exhibited greater disruption of capillary organization and reduced vascular integrity despite AAV8/BEST1-NRF2 infection (Fig. 6E, H-L). The choroidal vasculature was assessed using podocalyxin stained RPE-choroid flat mounts. In wild-type mice, AAV8/BEST1-NRF2 infection preserved a denser and more organized choriocapillaris network following NaIO3 injury. This choroidal protection was weakened in Gdf15⁻/⁻ animals, which exhibited lower capillary density and more pronounced vascular disorganization across posterior and peripheral regions despite NRF2 expression (Fig. 6F). Thus, GDF15 also contributes to the ability of AAV8/BEST1-NRF2 infection to preserve choroidal vascular structure under oxidative stress.

RPE morphology was evaluated by phalloidin staining to determine if GDF15 was required for AAV8/BEST1-NRF2 mediated protection following NaIO3 injury (Fig. 6G). In wild-type mice, AAV8/BEST1-NRF2 infection preserved the hexagonal RPE cellular mosaic after NaIO3 injury. In Gdf15⁻/⁻ mice, the RPE monolayer remained relatively preserved, although mild irregularities in cell boundaries were occasionally observed. Consistent with this observation of the RPE, cone counts were preserved by AAV8/BEST1-NRF2 in both wild-type and Gdf15⁻/⁻ mice, with no significant difference between NRF2-treated wild-type and Gdf15⁻/⁻ groups (Fig. S6). These findings suggest that GDF15 contributes only modestly to the protective effects of NRF2 on the RPE, and that other mechanisms are likely sufficient to protect the RPE from NaIO3 injury. Together, these findings indicate that loss of GDF15 partially attenuates NRF2-mediated protection of retinal function and vascular integrity.

### Pharmacological Inhibition of TGF-β Receptor Signaling Attenuates NRF2 Mediated Protection

To examine whether NRF2-mediated protection involves TGF-β-related signaling, TGF-β receptor signaling was pharmacologically inhibited by two inhibitors, SB431542 and LY364947, before oxidative injury (Fig. 7A). Wild-type mice received neonatal subretinal injection of AAV8/BEST1-NRF2 or AAV-CTL, followed by daily treatment with SB431542 and LY364947 beginning 3 days before NaIO3 administration in adulthood. FFA showed that when TGF-β receptor signaling was inhibited, eyes infected by AAV8/BEST1-NRF2 exhibited retinal hyperfluorescence after NaIO3 injury, comparable to that observed in AAV-CTL treated eyes (Fig. 7A, H). These findings suggest that TGF-β-related receptor signaling contributes to the vascular protective phenotype associated with NRF2 overexpression.

**Figure 7.**
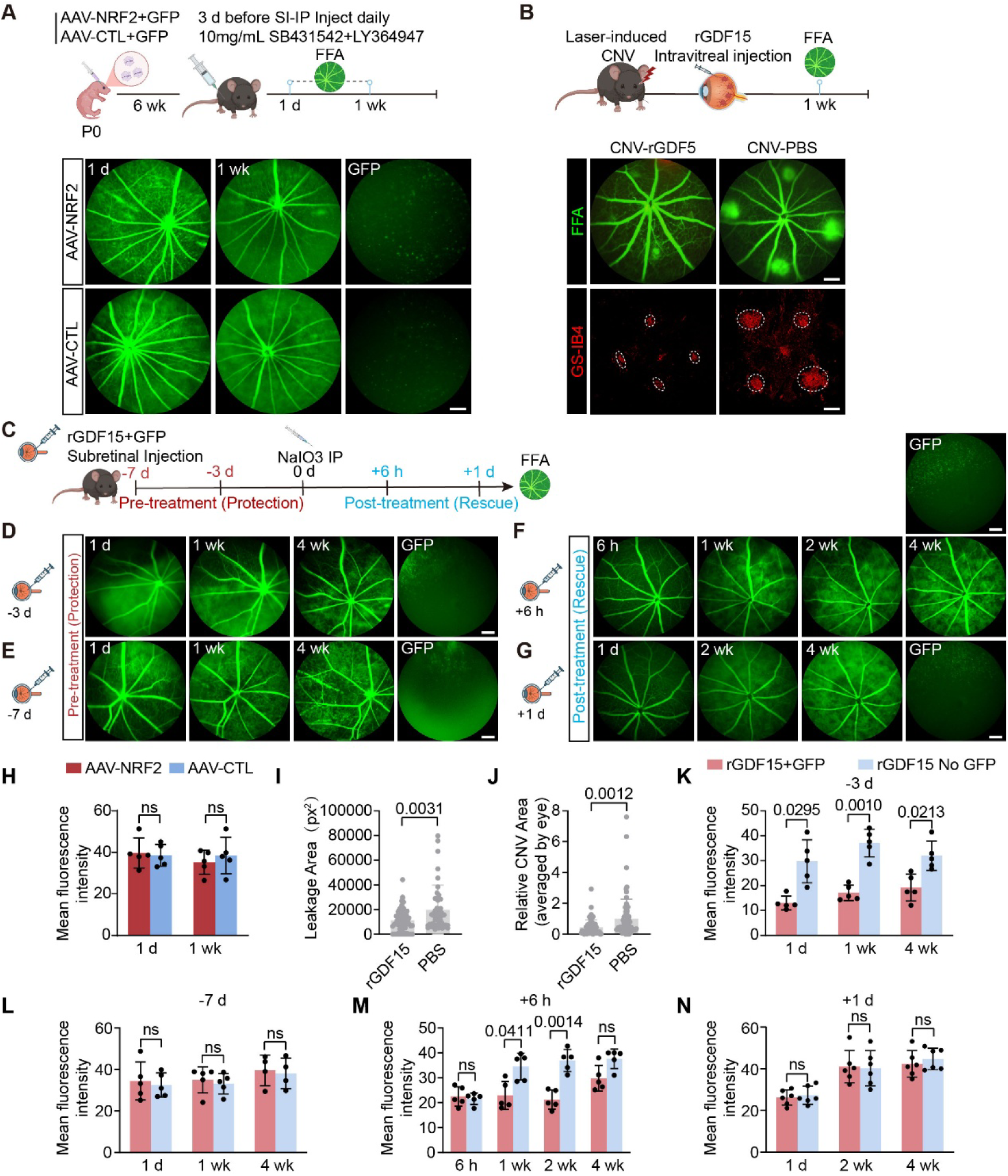
TGF-β pathway inhibition, test of CNV, and temporal analysis of GDF15-mediated protection in the NaIO3-injury model. **(A)** Experimental design for the inhibitor study. Wild-type mice received subretinal injection at P0 with AAV8/BEST1-NRF2 or AAV-CTL together with AAV-GFP. In adulthood, mice were treated daily intraperitoneal administration of SB431542 and LY364947, beginning 3 days before NaIO3 injection. FFA images over time are shown. Scale bar, 100 μm. **(B)** Experimental design for the laser-induced CNV model. FFA images and CNV lesion images are shown. Scale bar, 100 μm. **(C)** Experimental design for temporal analysis of GDF15 mediated protection. **(D-E)** Pretreatment analysis. Mice received subretinal injection of rGDF15 3 days **(D)**, 7days **(E)** before NaIO3. FFA images over time are shown. Scale bar, 100 μm. **(F-G)** Post-treatment analysis. Mice received rGDF15 6 hours **(F)**, 1 day**(G)** after NaIO3. FFA images over time are shown. Scale bar, 100 μm. **(H)** Quantification of fluorescein intensity shown in A (n = 5 eyes per group, mean ± SD, two-way ANOVA with Šídák’s multiple-comparisons test). **(I)** Quantification of leakage area of FFA shown in B. **(J)** Quantification of relative CNV lesion area shown in B (n = 19 eyes per group for PBS, n =17 eyes per group for rGDF15, mean ± SD, unpaired t-test). **(K–N)** Quantification of mean fluorescein intensity in GFP-positive and GFP-negative regions shown in D-G (n = 5-6 eyes per group, mean ± SD, two-way ANOVA with Šídák’s multiple-comparisons test).

### GDF15 Reduces CNV Lesion Size in a Laser-Induced CNV Model

To determine whether the vascular protective effects of GDF15 extend beyond NaIO3-induced oxidative injury, rGDF15 was tested in a laser-induced choroidal neovascularization (CNV) model (Fig. 7B). Intravitreal injection of rGDF15 was performed on the same day as laser injury, and FFA was carried out 7 days later to assess leakage of induced lesions. Representative FFA images and CNV flat mounts showed reduced hyperfluorescence and smaller CNV lesions in GDF15-treated eyes compared with controls (Fig. 7B). Quantification further confirmed that GDF15 treatment decreased CNV leakage and lesion size relative to controls (Fig. 7I-J). These findings indicate that GDF15 can reduce pathological vascular leakage and neovascular lesion development in the laser-induced CNV model.

### GDF15 Exerts Protection Within an Early Temporal Window

It was of interest to determine whether rGDF15 could confer protection when administered at different time points relative to oxidative injury (Fig. 7C). When rGDF15 was delivered 3 days before NaIO3 exposure, retinal vascular abnormalities were markedly reduced, (Fig. 7D, K). In contrast, when rGDF15 was administered 7 days before NaIO3 injury, no protective effect was observed (Fig. 7E, L). Shorter post injury time points also were tested. Administration of rGDF15 6 hours after NaIO3 injection partially reduced retinal hyperfluorescence (Fig. 7F, M), whereas administration 24 hours after injury showed minimal protective effect (Fig. 7G, N). These findings indicate that GDF15 mediated protection occurs within an early temporal window following oxidative injury.

## DISCUSSION

Oxidative stress is proposed to be a driver of degenerative change in dry AMD and related retinal disorders (1-3). The outer retina, RPE, Bruch’s membrane, and choriocapillaris function as a tightly coupled unit, so structural and molecular changes in one compartment may propagate to adjacent tissues (6-10). In the present study, systemic NaIO3 injury produced abnormalities in both retinal and choroidal vascular beds, with selective vulnerability of the DCP and marked disorganization and a reduced capillary density in the choriocapillaris. These changes were accompanied by substantial ERG deficits. Despite these acute and drastic changes, AAV8/BEST1-NRF2 was able to almost completely protect these tissues. The current results demonstrating the protection of vessels extends our earlier work showing that AAV8/BEST1-NRF2 overexpression preserves the RPE and photoreceptors, as well as vision, in the NaIO3 model as well as in models of retinitis pigmentosa (19, 20, 33).

A novel finding of this study is the demonstration that CM from NRF2 overexpressing RPE-choroid explants is sufficient to transfer protection in vivo. The ability of NRF2-CM to reduce abnormal angiographic signals, preserve deep plexus structure, and maintain choroidal vascular organization strongly support a non-cell-autonomous mechanism mediated by soluble factors, rather than by action only within cells that directly transcribe NRF2 target genes. The spatial restriction of protection to regions exposed to NRF2-CM suggests a local paracrine effect rather than a generalized systemic one. Our findings support a model in which NRF2 converts the stressed RPE from a source of injury propagation into a source of protective intercellular signaling. This finding suggests that RPE-centered therapies may protect vascular beds and neural tissue.

By integrating prior bulk RNA-seq data with single-cell receptor maps and validating the candidate at the protein and functional levels, we identified GDF15 as a downstream mediator of the NRF2-induced protection. GDF15 is a stress responsive cytokine with context dependent effects on metabolism, inflammation, and tissue adaptation (22, 23). In the eye, GDF15 is induced after retinal injury, can protect retinal ganglion cells, and can modulate retinal inflammation in autoimmune uveitis models (26-28). In the NaIO3 injury model, rGDF15 recapitulated major features of the NRF2-protective phenotype, whereas Gdf15-deletion partially weakened NRF2-mediated protection of vascular structure and retinal function. However, the phenotype was not completely eliminated in Gdf15-deficient mice, indicating that GDF15 should be regarded as an effector rather than the sole mediator of protection. NRF2 is a regulator of hundreds of genes and might be predicted to induce more than one protective factor (15).

The relationship between NRF2-CM and rGDF15 also supports the idea that GDF15 is an important but not exclusive component of the protective secretory response. Comparison of the amount of GDF15 in the CTL-CM and NRF2-CM revealed approximately a 2 fold difference, in keeping with the difference in the RNA-seq data (19). Dilution experiments with the CTL-CM and NRF2-CM revealed that there was more activity in the NRF2-CM than accounted for by GDF15 alone. In addition, the amount of GDF15 measured in the NRF2-CM was much lower than that required for protection by rGDF15. One interpretation is cooperativity: GDF15 may act together with other NRF2-induced soluble factors, thereby amplifying protection beyond what rGDF15 can achieve by itself. A second, not mutually exclusive, explanation is that endogenous GDF15 induced by NRF2 differs from rGDF15 in some way so as to increase its activity, perhaps by modifications that enhance its specific activity or stability.

Prior work in other systems has shown that endogenous GDF15 can participate in anti-inflammatory programs downstream of NRF2 activation e.g. (34, 35). Interestingly, rGDF15 was not able to recapitulate the effects predicted by GDF15 induction in hepatocytes by NRF2, leading to the suggestion that endogenous GDF15 may be more active due to modifications not made on recombinant forms (35).

The effects of GDF15 in this study were not restricted to vascular morphology. rGDF15 also preserved the RPE and sustained more photoreceptors than the control after injury. It is not clear if the effects of rGDF15 are via one particular cell type, with indirect effects on others, or whether more than one cell type responds to the addition of rGDF15. One interesting aspect of the effects of GDF15 on the RPE concerns its sufficiency vs. necessity. Addition of rGDF15 was sufficient to protect the RPE. However, AAV8/BEST1-NRF2 still protected the RPE in the Gdf15^-/-^ mice. Protection of the RPE in the absence of GDF15 may not be surprising given that NRF2 induces many more genes than GDF15 in the RPE (19), likely at least providing protection from oxidative damage. This makes the sufficiency of rGDF15’s protection of the RPE all the more intriguing.

The protection of vessels by either NRF2 overexpression in the RPE, or by addition of GDF15, can be considered in light of AMD pathology. Wet AMD occurs when choroidal vessels invade the retina, perhaps in response to inadequate trophic support from an ailing RPE (36). By improving the health of the RPE, potentially in dry AMD, the probability of conversion of dry to wet AMD could be reduced. Similarly, the fairly broad protection of vessels, RPE and photoreceptors may predict a therapeutic effect of NRF2 overexpression in the RPE, or addition of GDF15, for diabetic retinopathy where vessels are affected (5).

Mechanistically, the receptor landscape for GDF15 in peripheral tissues remains incompletely resolved. While circulating GDF15 signals through the hindbrain GFRAL receptor, which recruits RET for its effects (24, 25), GFRAL expression has been documented to be absent in peripheral tissues and outside of the brainstem in the brain (37). Our ligand-receptor analysis and TGF-β receptor inhibitor experiments are consistent with signaling through the TGF-β receptor, and SMAD phosphorylation following addition of GDF15 to retinal cultures is consistent with this signaling pathway (38). However, additional work will be needed to establish TGF-β receptor as the relevant receptor. It is also worth noting that the rGDF15 used here was produced in E. coli, making contamination by mammalian TGF-β family proteins during production a less likely explanation for the observed activity (39).

The laser-induced CNV model is widely used as an experimental surrogate for neovascular AMD (40, 41), The CNV experiments conducted here show that the protective activity of GDF15 is not restricted to the NaIO3 model. When rGDF15 was delivered intravitreally on the day of laser injury, the injured eyes showed reduced fluorescein leakage at day 7, together with smaller CNV lesions. Additionally, the protection of ganglion cells, amelioration of uveitis, as well as the activities of GDF15 in other disease models, suggest that GDF15 might not only be a marker of stress, but can exert protective effects more broadly (26, 27).

Finally, the temporal dependence of protection is one of the most clinically informative aspects of this work. Protection was evident when rGDF15 was delivered 2-3 days before injury and was still effective when administered 6 hours after NaIO3. However, protection was lost with a 7 day pretreatment interval or a 1 day post injury delay. These findings indicate both a limited preconditioning window and a short early phase during which injury remains biologically reversible. Oxidative injury likely initiates rapidly, but the downstream collapse of vascular stability, barrier integrity, and inflammatory homeostasis likely unfolds over time. Once that cascade progresses beyond a threshold, later intervention becomes ineffective. This interpretation is consistent with broader work in oxidative retinal degeneration showing that timing strongly constrains rescue efficacy (3, 13).

## MATERIALS AND METHODS

### Animals

Adult C57BL/6J mice were used for wild-type experiments. Timed pregnant females were used for neonatal subretinal injections. Gdf15 knockout mice were obtained from Dr. Randy Seeley and maintained as homozygous breeding pairs (42). Both male and female mice were included. All procedures were approved by the Harvard University Institutional Animal Care and Use Committee.

### AAV Vectors and Subretinal Delivery

AAV8/BEST1-NRF2 was used to express human NRF2 in RPE cells. AAV-CTL served as the control vector, and AAV-GFP was co-administered to mark transduced regions (20). Vectors were produced in HEK293T cells and purified by iodixanol gradient ultracentrifugation (33).

For neonatal injections, P0 pups received subretinal AAV8/BEST1-NRF2 or AAV-CTL at 4 × 10^8 vector genomes (vg)/eye plus AAV-GFP tracer at 1 × 10^7 vg/eye (20). Approximately 0.25 μL was delivered per eye. One eye received AAV8/BEST1-NRF2 and the contralateral eye received AAV-CTL, with laterality alternated across litters. Animals with less than 80% posterior RPE transduction were excluded.

For adult injections, 6-8-week-old mice received approximately 1 μL subretinal viral suspension. AAV8/BEST1-NRF2 or AAV-CTL was administered at 1 × 10^9 vg/eye with AAV-GFP tracer. Transduction was typically limited to the injection bleb region.

### Conditioned Medium Preparation, Administration, and ELISA

For CM experiments, mice received P0 subretinal AAV8/BEST1-NRF2 or AAV-CTL injections. At 8 weeks, RPE-choroid complexes were dissected and cultured RPE side up for 24 h in serum-free explant medium with three explants per 500 μL. For subretinal administration, centrifuged media were separated using 10-kDa molecular-weight cutoff filters, and adult recipient mice received 1 μL of the indicated NRF2-CM or CTL-CM fraction with GFP tracer, followed by NaIO3 injury two days later. For dilution experiments, centrifuged media were diluted 1:10 before injection. GDF15 levels were measured separately using uncentrifuged CM with the Mouse/Rat GDF-15 Quantikine ELISA Kit.

### Recombinant Protein Administration

Recombinant mouse GDF15 was administered subretinally to adult mice, typically two days before NaIO3 injury. Unless otherwise stated, rGDF15 was delivered at 10 μM in 1 μL per eye (28). Dose-response experiments used 50, 10, or 1 ng per eye. For timing experiments, rGDF15 was injected 7, 3, or 2 days before, or 6 h or 1 day after, NaIO3 injury. The contralateral eye received PBS. Recombinant mouse oncostatin M was delivered subretinally at 100 ng per eye (43).

### NaIO3 Injury and CNV Model

NaIO3 was freshly dissolved in sterile saline and administered intraperitoneally at 75 mg/kg. Control mice received saline. For laser-induced CNV, four laser burns were applied per eye using the Micron IV image-guided system with a 532-nm laser (44). Burns were placed equidistant from the optic nerve, and only lesions with a vaporization bubble were included. Immediately after laser injury, mice received intravitreal rGDF15 or PBS. At 7 days, FFA was performed and eyes were collected for flat-mount analysis.

### FFA and ERG

FFA was performed using the Micron IV system. After intravenous injection of 1% sodium fluorescein, images were acquired 1 min later. In the NaIO3 model, angiographic hyperfluorescence was quantified in ImageJ by measuring extravascular fluorescence while excluding major retinal vessels. For adult partial-transduction experiments, analysis was restricted to GFP-positive regions. For CNV, individual lesions were manually outlined and hyperfluorescent lesion area was quantified.

Ganzfeld ERG was performed using an Espion E3 system (20). Mice were dark-adapted for at least 2 h before recording. Scotopic responses, oscillatory potentials, and photopic responses after light adaptation were collected and analyzed using Espion software.

### Pharmacological Inhibition

To inhibit TGF-β receptor signaling, mice received daily intraperitoneal SB431542 and LY364947 at 10 mg/kg each, beginning 3 days before NaIO3 injection and continuing until 7 days after injury (45).

### Tissue Processing, Staining, and Imaging

Eyes were dissected into neural retina and RPE-choroid complexes. Retinas were stained with GS-IB4 to label retinal vasculature or with anti-cone arrestin antibody for cone analysis. RPE flat-mounts were stained with phalloidin to visualize RPE cell morphology. For choroidal vascular staining, RPE-choroid tissues were treated with EDTA to remove RPE, bleached with hydrogen peroxide, and stained with anti-podocalyxin antibody (46). Samples were mounted as flat-mounts and imaged using an Olympus VS200 slide scanner or fluorescence imaging systems.

### Image Analysis

Image analysis was performed using ImageJ/Fiji and REAVER. For RPE quantification, four regions were sampled from each flat-mount, avoiding damaged or out-of-focus areas, and RPE cells were manually counted (20). For adult partial-transduction experiments, GFP-positive and GFP-negative regions were analyzed separately. Retinal vascular parameters were quantified using REAVER (47), including vessel area fraction, vessel length, segment count, and branchpoint number. Cone numbers were manually counted from representative regions.

### Statistics

Statistical analyses were performed using GraphPad Prism and Microsoft Excel. Unless otherwise stated, n represents individual eyes. Data are presented as mean ± SD. Paired or unpaired two-tailed Student’s t tests, nonparametric tests, one-way ANOVA, or two-way ANOVA with multiple-comparisons correction were used as appropriate. Contralateral-eye comparisons were analyzed as paired samples, and GFP-positive versus GFP-negative regions within the same eye were analyzed using eye-level paired summary values.

## Acknowledgments

We thank Dr. Randy Seeley (University of Michigan) for generously providing the Gdf15 knockout mice. This study was supported by the Howard Hughes Medical Institute.

## Author Contributions

S.W., and C.L.C., designed research; S.W., S.R.Z., A.D., A.G., E.N., and X.T. performed research, S.W., A.D., and C.L.C. analyzed the data. S.W. and C.L.C. wrote the paper with feedback and support from L.S. and Z.F.

## Significance Statement

Age-related macular degeneration is a multicompartment disease involving the retinal pigment epithelium, photoreceptors, and ocular vasculature. How protective signals might coordinate across these tissues remains unclear. Using a mouse model of oxidative injury, we show that overexpressing NRF2 only in the retinal pigment epithelium preserves retinal and choroidal vascular structure. We identify GDF15 as a functionally important downstream mediator. Recombinant GDF15 reproduces major protective effects in vivo, extending beyond the vasculature to preservation of RPE and photoreceptors, and Gdf15-deficiency weakens NRF2-mediated protection. These findings indicate that NRF2 can induce the RPE to produce factors that protect vasculature as well as photoreceptors and identify GDF15 as a candidate therapeutic for AMD and other disorders with vascular pathology.

