## Supplementary material for "AAV-NRF2 protects retinal and choroidal vasculature in a GDF15-dependent manner in an oxidative damage model of AMD": Supplymentary Method

**MATERIALS AND METHODS**

**Animals**

Adult C57BL/6J mice (The Jackson Laboratory, strain 000664) were used for wild-type experiments. Timed pregnant females were purchased for neonatal subretinal injection experiments. Gdf15 knockout (Gdf15**⁻/⁻**) mice were originally generated by CRISPR/Cas9-mediated deletion of exon 2 of the Gdf15 gene and were obtained from Dr. Randy Seeley (University of Michigan) (1). Gdf15**⁻/⁻** neonatal (P0) pups used in this study were derived from homozygous breeding pairs maintained in house. Both male and female animals were included. All procedures were approved by the Harvard University Institutional Animal Care and Use Committee (IACUC protocol #1695), and animals were maintained under a 12-h light/dark cycle with ad libitum access to food and water.

**AAV Vector Design and Production**

An AAV8 vector expressing human NRF2 under the control of the RPE-specific BEST1 promoter (AAV8/BEST1-NRF2) was used for targeted gene delivery to the retinal pigment epithelium (RPE). A transcriptionally inactive control vector containing a 6×STOP cassette (AAV8/BEST1-6xSTOPmutGFP; hereafter AAV-CTL) was used as a matched control (2). A nuclear GFP tracer vector (AAV8/RedO-H2B-GFP; hereafter AAV-GFP) was co-administered where indicated to demarcate the transduced region. Vectors were produced in house by transient transfection of HEK293T cells and purified by iodixanol gradient ultracentrifugation, as previously described (3). Vector genome titers were determined by quantitative comparison with reference AAV stocks of known concentration.

**Neonatal Subretinal Injections**

Both wild-type and Gdf15⁻/⁻ P0 pups were anesthetized by brief hypothermia and received buprenorphine (0.05-0.1 mg/kg, subcutaneous) immediately before injection. The palpebral fissure was opened with a 30-gauge needle, and a hand-pulled beveled glass micropipette was advanced into the subretinal space under direct visualization, as described previously (2). AAV8/BEST1-NRF2 or AAV-CTL was delivered at 4 × 10^8 vg/eye, together with AAV-GFP at 1 × 10^7 vg/eye. One eye received AAV8/BEST1-NRF2 and the contralateral eye received AAV-CTL, with injection laterality alternated across litters to minimize bias. Approximately 0.25 μL was delivered per eye, and successful injections were confirmed by formation of a visible subretinal bleb. A drop of 0.5% proparacaine hydrochloride ophthalmic solution was applied after injection. Animals with less than 80% RPE transduction in the posterior pole were excluded before statistical analysis.

**Adult Subretinal Injections**

Adult wild-type mice (6–8 weeks of age) were anesthetized with ketamine (80 mg/kg) and xylazine (10 mg/kg) by intraperitoneal injection. A small incision was made at the corneoscleral junction, and a beveled glass micropipette was inserted into the subretinal space. Approximately 1 μL of viral suspension was delivered per eye to generate a localized retinal detachment. AAV8/BEST1-NRF2 or AAV-CTL was administered at 1 × 10^9 vg/eye, together with AAV-GFP at 1 × 10^7 vg/eye. In adult mice, transduction was typically restricted to the injection bleb region. For flat-mount analyses, GFP-positive and GFP-negative regions were quantified separately where indicated. Injection laterality was alternated across cohorts to minimize bias.

**Preparation and Subretinal Administration of NRF2-CM**

Wild-type mice received subretinal injection of AAV-NRF2 or AAV-CTL at P0 as described above. At 6–8 weeks of age, mice were euthanized and RPE-choroid complexes were immediately dissected under a stereomicroscope. Explants were cultured RPE side up in explant medium (DMEM supplemented with GlutaMAX, MEM non-essential amino acids, and penicillin-streptomycin; all supplied as 100× stock solutions) for 24 h at 37 °C, with 3 explants per 500 μL of medium. Culture media were collected, cleared by centrifugation, and fractionated using 10 kDa molecular-weight cutoff filters (Amicon Ultra-0.5; MilliporeSigma, UFC5010). For subretinal delivery, 1 μL of NRF2-conditioned medium (NRF2-CM) or control-conditioned medium (CTL-CM) was injected into adult wild-type mice together with the AAV-GFP tracer. Both eyes were injected in each animal, with treatment assignment alternated between left and right eyes to minimize lateralization bias. Two days later, mice received NaIO3 as described below. For dilution experiments, conditioned media were diluted 1:10 in serum-free medium before injection.

**Recombinant GDF15 Administration**

Recombinant mouse GDF15 (rGDF15; R&D Systems, #8944-GD-025) was administered to adult wild-type mice by subretinal injection using the same surgical approach described above. Unless otherwise indicated, rGDF15 was delivered at 10 μM in a total volume of 1 μL per eye (4), typically 2 days prior to NaIO3 administration. For dose-response experiments, rGDF15 was administered subretinally at 50, 10, or 1 ng per eye in a total volume of 1 μL. Both eyes were injected in each animal, with one eye receiving rGDF15 and the contralateral eye receiving PBS. AAV-GFP tracer vector was co-administered as described above. Assignment of treatment to the left or right eye was alternated across animals to minimize lateralization bias. GFP fluorescence was used to define the injected region for downstream analyses. For temporal window experiments, rGDF15 was administered at different time points relative to NaIO3 injection, including 7, 3, or 2 days before, or 6 hours or 1 day after, NaIO3 administration.

For the laser-induced choroidal neovascularization (CNV) model, immediately after laser photocoagulation, mice received bilateral intravitreal injection of either rGDF15 at 10 μM in 1 μL per eye or PBS at 1 μL per eye using a Hamilton syringe. Eyes were rinsed with sterile saline, treated with erythromycin ophthalmic ointment (Fougera), and maintained on a 35 °C heating pad until recovery. The 10 μM dose was selected with reference to prior studies using intraocular rGDF15 in ocular disease models (5), whereas lower doses were tested in dose-response experiments.

**Recombinant OSM Administration**

Recombinant mouse oncostatin M (OSM; R&D Systems, #495-MO-025) was administered to adult wild-type mice by subretinal injection using the same surgical approach described above. OSM was delivered at 100 ng per eye in a total volume of 1 μL, typically 2 days prior to NaIO3 administration. For each animal, one eye received OSM and the contralateral eye received PBS as a control. AAV-GFP tracer was co-administered as described above. The dose was selected with reference to prior studies using intraocular OSM in retinal disease models (6).

**Sodium Iodate-Induced Oxidative Injury**

Sodium iodate (NaIO3) was freshly dissolved in sterile saline and administered by intraperitoneal injection at 75 mg/kg to induce acute oxidative injury. Control animals received an equivalent volume of saline.

**Fluorescein Angiography and Analysis**

Fundus fluorescein angiography (FFA) was performed using the Micron IV Retinal Imaging Microscope system (Phoenix Research Labs), as previously described for sodium iodate injury models (7). Following anesthesia and pupil dilation, 1% sodium fluorescein was administered intravenously. Fundus images were acquired 1 minute after injection. For the NaIO3 model, the fluorescein intensity was quantified by measuring extravascular fluorescence area and intensity using ImageJ (NIH) in a masked manner. Major retinal vessels were excluded from analysis to avoid confounding intravascular signal. For spatial analyses, GFP-positive and GFP-negative regions were quantified separately within the same retina. For the laser-induced CNV model, FFA images were analyzed in a masked manner using ImageJ. Individual CNV lesions were manually outlined as regions of interest, and leakage was quantified by measuring the hyperfluorescent lesion area.

**Electroretinography (ERG)**

Ganzfeld ERG was performed using an Espion E3 system (Diagnosys LLC), and data were analyzed with Espion V6 software. Mice were dark-adapted for at least 2 h before recording. Scotopic ERG was recorded at 0.1 cd·s/m², followed by oscillatory potentials at 3 cd·s/m². After 6 min of light adaptation, photopic responses were recorded at 1, 10, 100, and 1000 cd·s/m² on a 30 cd·s/m² white background (2).

**Retina and RPE-Choroid Dissection**

Mouse eyes were enucleated and the anterior segment, including the cornea and lens, was removed under a dissecting microscope. The neural retina was carefully separated from the underlying RPE-choroid complex. For experiments shown in Fig. 6 using Gdf15**⁻/⁻** mice, the RPE-choroid complex was bisected through the central region, with one half used for RPE flat-mount staining and the other half processed for choroidal vascular staining.

**Retinal Flat-Mount Staining**

Retinas were fixed in 4% paraformaldehyde for 4 h at room temperature and washed three times in PBS. Tissues were permeabilized in 0.5% Triton X-100 in PBS overnight. For visualization of retinal vasculature, retinas were incubated with Isolectin GS-IB4 (Invitrogen, #121413) at 20 μg/mL for 2 h at room temperature. For cone staining, retinas were incubated with rabbit anti-cone arrestin antibody (MilliporeSigma, #AB15282,1:2000) overnight at 4 °C. Samples were washed three times in PBS and incubated with Alexa Fluor 647-conjugated donkey anti-rabbit secondary antibody (Jackson ImmunoResearch, #711-605-152) at 1:750 dilution in PBS for 1 h at room temperature. After staining, retinas were washed three times in PBS and flat-mounted into four petals, with the ganglion cell layer facing the coverslip.

**RPE Flat-Mount Staining**

After removal of the neural retina, the RPE-choroid complex was fixed in 4% paraformaldehyde for 4 h at room temperature and washed three times in PBS. Tissues were incubated overnight at 4 °C with Alexa Fluor 568 phalloidin (1:100 dilution in staining buffer) to visualize the actin cytoskeleton of RPE cells. After staining, tissues were washed three times in PBS and flat-mounted into eight petals with the RPE surface facing the coverslip.

**Choroidal Vascular Staining**

To visualize choroidal vasculature, the RPE-choroid complex was incubated in 1% EDTA in PBS for 2 h at room temperature to facilitate removal of the RPE layer. Samples were washed three times in PBS and fixed in 2% paraformaldehyde overnight at 4 °C. Tissues were then washed three times in PBS containing 0.1% Triton X-100. To reduce pigmentation, samples were incubated in 10% hydrogen peroxide in PBS at 55 °C for 90 minutes. After bleaching, tissues were washed three times in PBS and incubated overnight at 4 °C with rat anti-mouse podocalyxin antibody (R&D Systems, #MBA1556) at 1:100 dilution to label choroidal endothelial cells. Samples were washed three times in PBS and incubated with Alexa Fluor 647-conjugated donkey anti-rat secondary antibody (Jackson ImmunoResearch, #712-605-153) at 1:750 dilution in PBS for 1 h at room temperature. After secondary incubation, tissues were washed three times in PBS before mounting (8).

**Retinal Cryosectioning**

For cross-sectional analysis, previously imaged retinal flat-mount samples were carefully removed from slides and transferred to a cryoprotection solution consisting of a 1:1 mixture of Optimal Cutting Temperature compound and 30% sucrose for at least 30 minutes at 4 °C. Samples were embedded in OCT compound and cryosectioned at 20 μm thickness. Sections were stained with DAPI for at least 15 minutes prior to imaging.

**Pharmacological Inhibition of TGF-β Receptors**

To inhibit TGF-β receptor signaling in vivo, mice were treated with a combination of SB431542 (SelleckChem, #S1067) and LY364947 (SelleckChem, #S2805). Both compounds were dissolved in PBS containing 5% dimethyl sulfoxide (DMSO) and 30% polyethylene glycol 300. Mice received daily intraperitoneal injections at 10 mg/kg for each compound. Drug administration was initiated 3 days prior to NaIO3 injection and continued until 7 days after NaIO3 administration. This inhibitor-based approach was informed by prior work on TGF-β signaling in retinal degeneration (9).

**Quantification of GDF15 in Conditioned Medium by ELISA**

GDF15 levels in conditioned medium were measured using a Mouse/Rat GDF-15 Quantikine ELISA Kit (R&D Systems, MGD150) according to the manufacturer’s instructions. Unfractionated conditioned medium from AAV8-BEST1-NRF2 and control RPE-choroid explants was analyzed directly without filtration or molecular-weight separation. After brief centrifugation to remove debris, samples were assayed using 50 µL per well, and absorbance was measured at 450 nm. Concentrations were interpolated from the standard curve and compared across independent biological replicates.

**Laser-Induced Choroidal Neovascularization Model**

Mice were anesthetized, pupils were dilated with Cyclomydril (Alcon Laboratories), and corneas were kept hydrated with lubricating eye drops. Laser-induced choroidal neovascularization (CNV) was generated as previously described using an image-guided laser system (10). Using the Micron IV image-guided platform (Phoenix Research Laboratories), four laser burns were applied per eye at positions equidistant from the optic nerve with a 532-nm laser (50 μm spot size, 70 ms duration, 300 mW power). Only burns associated with the appearance of a vaporization bubble were considered successful. Immediately after laser photocoagulation, mice received bilateral intravitreal injection of either rGDF15 or PBS, as described above. Eyes were rinsed with sterile saline, treated with erythromycin ophthalmic ointment (Fougera), and mice were maintained on a 35 °C heating pad until recovery. At 7 days after laser injury, FFA was performed to assess CNV-associated leakage, and eyes were subsequently collected for flat-mount analysis. For CNV lesion quantification, RPE-choroid-sclera flat-mounts were prepared and stained with GS-IB4. CNV lesion areas were measured from fluorescence images using ImageJ, and the lesion area from each burn was quantified individually. Lesions with obvious hemorrhage, fused lesions, or unsuccessful burns were excluded from analysis.

**Imaging and Image Analysis**

RPE flat-mounts and retinal sections were imaged using an Olympus VS200 slide scanner. Retinal flat-mounts were imaged using a fluorescence lifetime imaging microscopy (FLIM) system equipped with a white-light laser and pulse picker. Images were processed and analyzed using ImageJ/Fiji software.

For RPE flat-mount quantification, a custom Fiji macro was used to extract four equally sized regions (~0.078 mm² each) from full-sized phalloidin-channel images. Region placement was determined by lines drawn by the investigator from the outer RPE boundary to the optic nerve head (ONH). For NaIO3-treated eyes, regions were selected within the GFP-positive transduced area, whereas in saline-treated controls, regions could be selected anywhere on the flat-mount. The midpoint of each line was used to define one corner of each sampling box. Care was taken to distribute boxes approximately evenly and to avoid out-of-focus or damaged regions. RPE cells within each region were manually counted using the Fiji Cell Counter function, and the median value from the four sampled regions was reported. In occasional cases, suboptimal regions were excluded from the median calculation. For adult eyes with partial transduction, the macro was applied separately to transduced and untransduced regions defined by GFP signal, and median RPE counts were calculated for each region independently. For control vector-injected RPE flat-mounts with central cell loss after NaIO3 treatment, transduced regions were defined by surviving peripheral H2B-GFP-positive RPE cells near the ciliary margin (2).

For retinal flat-mount vascular analysis, tissues were stained with GS-IB4 to visualize the vascular network. Two representative rectangular regions (~500 × 500 μm) were extracted from each full-sized retinal flat-mount image, avoiding out-of-focus areas or regions with tissue damage. GFP-positive and GFP-negative regions were sampled separately where applicable. Extracted images were analyzed using the Rapid Editable Analysis of Vessel Elements Routine (REAVER), an open-source MATLAB-based platform for vascular image quantification (11). The following parameters were measured: vessel area fraction, total vessel length, segment count, branchpoint number, and maximal diffusion distance. The mean value from the two sampled regions was reported.

For cone survival analysis, representative rectangular regions of defined size (500 × 500 μm) containing labeled cones were extracted from full-sized retinal flat-mount images. Regions were selected within GFP-positive areas when a AAV-GFP tracer was used. Following the general sampling strategy used in Gardner et al (2), regions were chosen from the transduced retinal area while avoiding poor focus, weak staining, tissue-processing artifacts, or obvious damage. Cone numbers within each extracted region were manually counted by the investigator using ImageJ/Fiji.

FFA images were quantified by excluding major vascular structures and analyzing the remaining background fluorescence as a measure of angiographic hyperfluorescence (12). For adult mouse subretinal injection experiments with partial transduction, the quantified region was restricted to the GFP-positive area by deleting non-transduced regions in ImageJ/Fiji. Mean fluorescence intensity of the remaining background area was then calculated. For fundus angiograms from CNV experiments, total CNV lesion area was quantified directly using ImageJ/Fiji.

**Sample Exclusion Criteria**

Flat-mount data from both eyes were excluded when there was evidence of failed intraperitoneal injection or a general lack of NaIO3-induced damage in the control AAV/NaIO3 eye. Individual retina, RPE, or section datasets were excluded if tissues were too damaged during dissection to permit reliable flat-mounting or quantification, or if antibody staining, immunohistochemistry, or imaging artifacts prevented analysis. For NaIO3-treated mice, retinal flat-mounts from NRF2-treated eyes were excluded when RedO-H2B-GFP indicated incomplete transduction. For two mouse ERG cohorts, ERG data were excluded when histology indicated incomplete transduction of the NRF2-treated eye or when no histological reference data were available.

**Statistics**

All statistical analyses were performed using GraphPad Prism version 9.5.0 and Microsoft Excel. Unless otherwise indicated, n represents individual eyes. A P value < 0.05 was considered statistically significant, and P ≥ 0.05 was considered not significant (ns). Group data are presented as mean ± SD unless otherwise indicated.

Statistical tests were selected according to the experimental design and the distribution of the data. For comparisons between two groups, unpaired or paired two-tailed Student’s t tests were used as appropriate. For paired analyses, only complete pairs were included. For datasets that did not satisfy assumptions for parametric testing, or for which normality could not be reliably assessed, nonparametric tests were used as appropriate. When multiple pairwise comparisons were performed within the same experiment, multiple-comparisons correction was applied as indicated in the figure legends. For comparisons among three or more groups, one-way ANOVA with an appropriate multiple-comparisons test was used. For experiments involving two independent variables, two-way ANOVA with Šídák’s multiple-comparisons test was used where appropriate.

For contralateral-eye experiments, paired analyses were performed by eye, with the two eyes from the same animal treated as matched samples. For comparisons of GFP-positive and GFP-negative regions within the same eye, paired statistical tests were performed using eye-level summary values generated separately for each region.
