## Supplementary material for "AAV-NRF2 protects retinal and choroidal vasculature in a GDF15-dependent manner in an oxidative damage model of AMD": Supplymentary figures and legends

**Supplementary Figures and Legends**

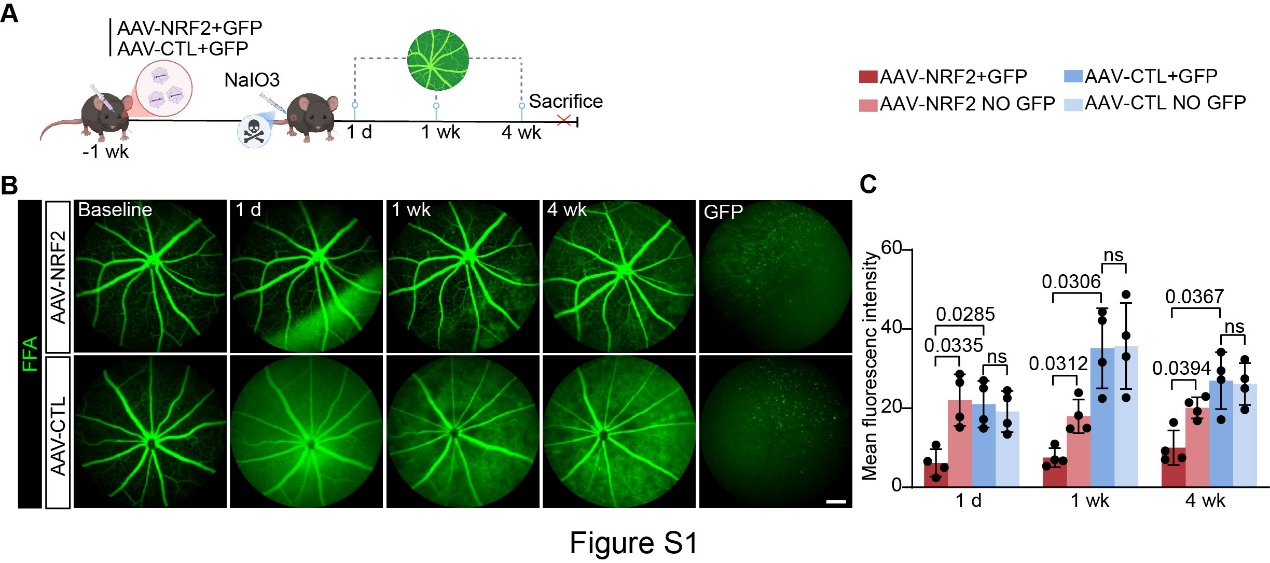

**Figure S1. Adult subretinal AAV8/BEST1-NRF2 delivery reduces NaIO3-induced angiographic abnormalities.**

**(A)** Adult mice were subretinally injected with AAV8/BEST1-NRF2 or AAV8/BEST1-6xSTOPmutGFP control vector (AAV-CTL), together with a AAV-GFP tracer vector. After 1 week, mice were treated with NaIO3, and fundus fluorescein angiography (FFA) was performed at baseline, 1 day, 1 week, and 4 weeks after injury.

**(B)** FFA images and corresponding GFP fundus images are shown. Scale bar, 100 μm.

**(C)** Quantification of mean fluorescein intensity from the GFP-positive regions shown in B. GFP images were used to identify transduced regions in eyes co-injected with the tracer vector (n = 4 eyes per group, mean ± SD, two-way ANOVA with Tukey’s multiple-comparisons test).

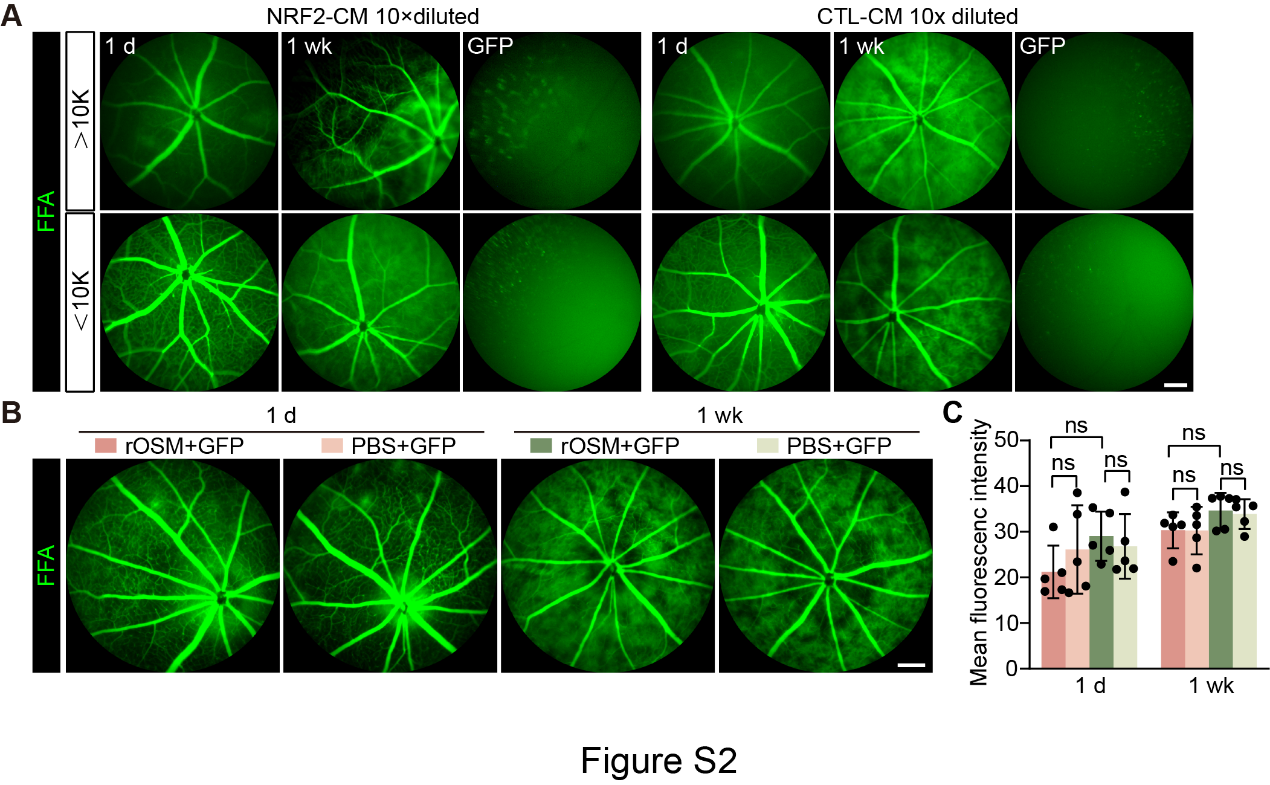

**Figure S2. Dilution sensitivity of conditioned medium fractions and testing of recombinant oncostatin M after NaIO3 injury.**

**(A)** FFA images at 1 day after NaIO3 administration following subretinal injection of 10× diluted >10 kDa or <10 kDa conditioned medium (CM) fractions. Scale bar, 100 μm.

**(B)** FFA images at 1 day and 1 week after NaIO3 administration in eyes receiving subretinal injection of recombinant oncostatin M (OSM) or PBS. Scale bar, 100 μm.

**(C)** Quantification of mean fluorescein intensity shown in B (n = 5 eyes per group, mean ± SD, two-way ANOVA with Tukey’s multiple-comparisons test).

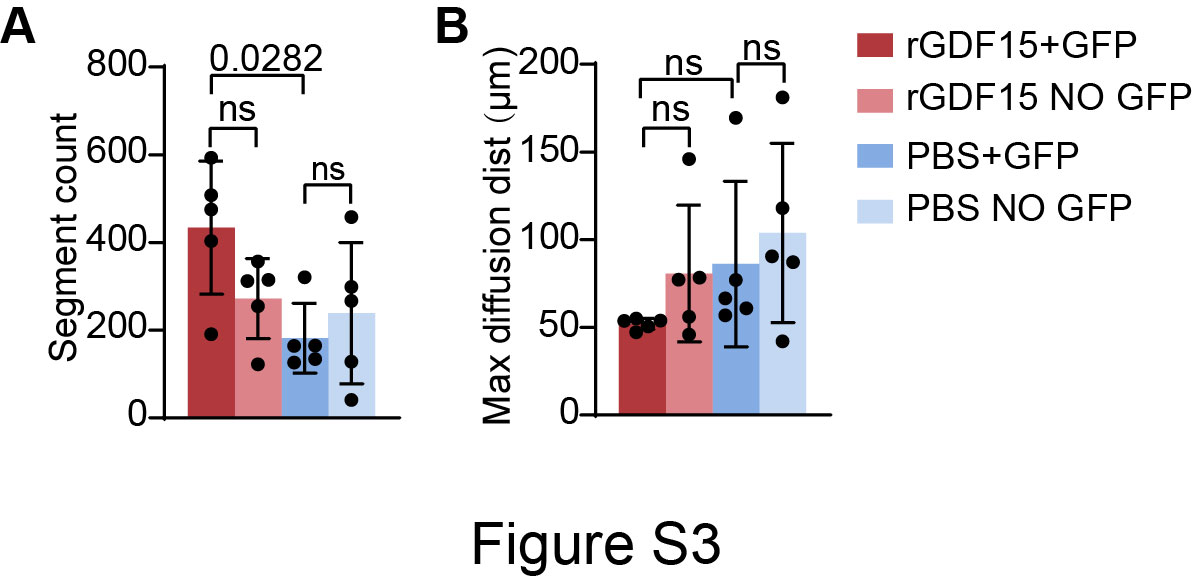

**Figure S3. Recombinant GDF15 partially preserves retinal vascular architecture.**

**(A-B)** Quantification of retinal vascular parameters in GFP-positive and GFP-negative regions of recombinant GDF15 (rGDF15) - or PBS-treated eyes after NaIO3 injury. Segment count **(A)** and maximal diffusion distance **(B)** are shown for the deep capillary plexus (DCP) images presented in Figure 5C (n = 5 eyes per group, mean ± SD, one-way ANOVA with Tukey’s multiple-comparisons correction).

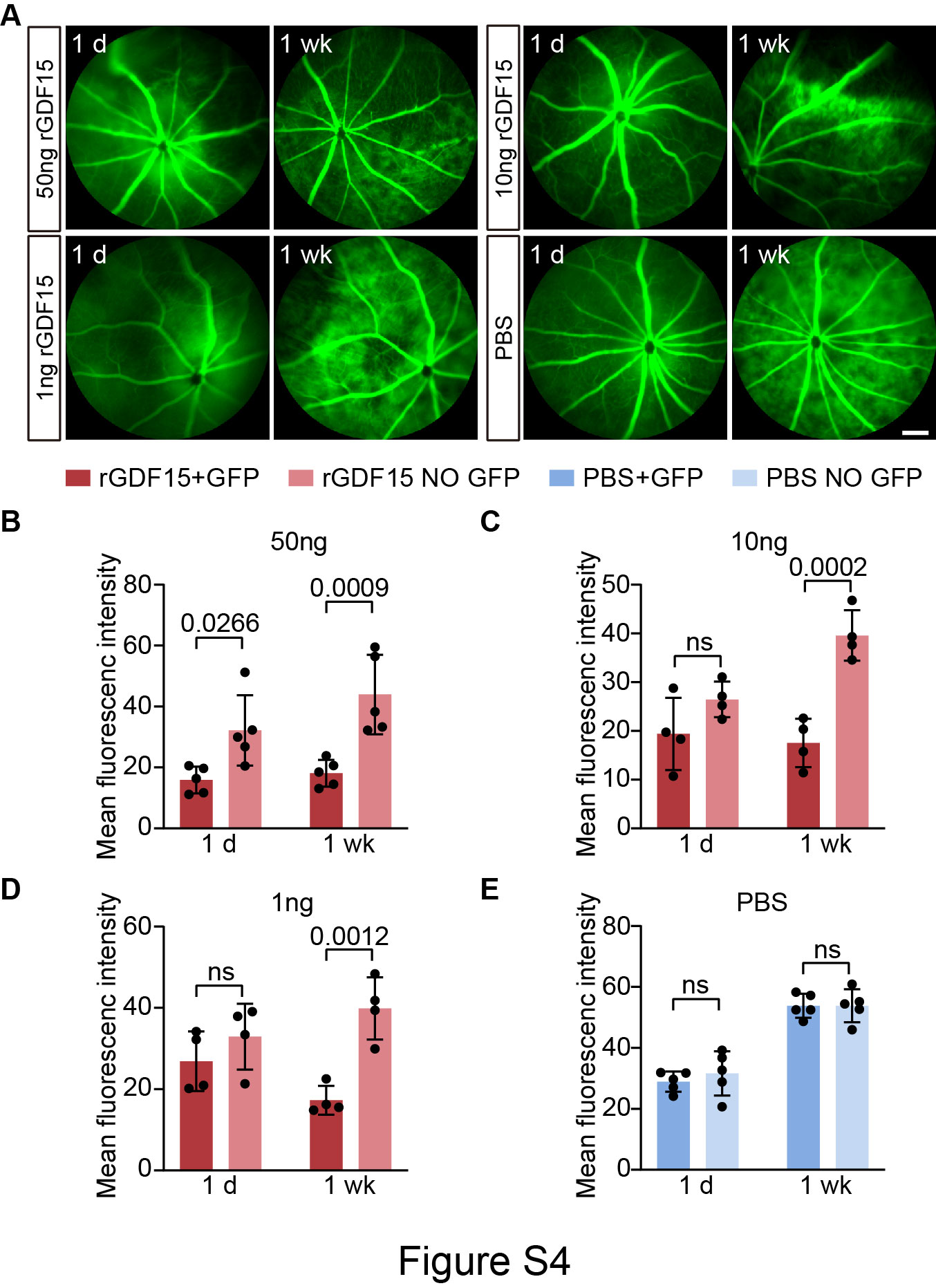

**Figure S4. Dose-dependent and time-dependent effects of recombinant GDF15 after NaIO3 injury.**

**(A)** FFA images from eyes receiving subretinal injection of rGDF15 at 50 ng, 10 ng, or 1 ng, or PBS, together with GFP tracer. Images are shown at 1 day and 1 week after NaIO3 administration. Scale bar, 100 μm.

**(B–E)** Quantification of mean fluorescein intensity in GFP-positive and GFP-negative regions of rGDF15 or PBS-treated groups (n = 4-5 eyes per group, mean ± SD, two-way ANOVA with Šídák’s multiple-comparisons test).

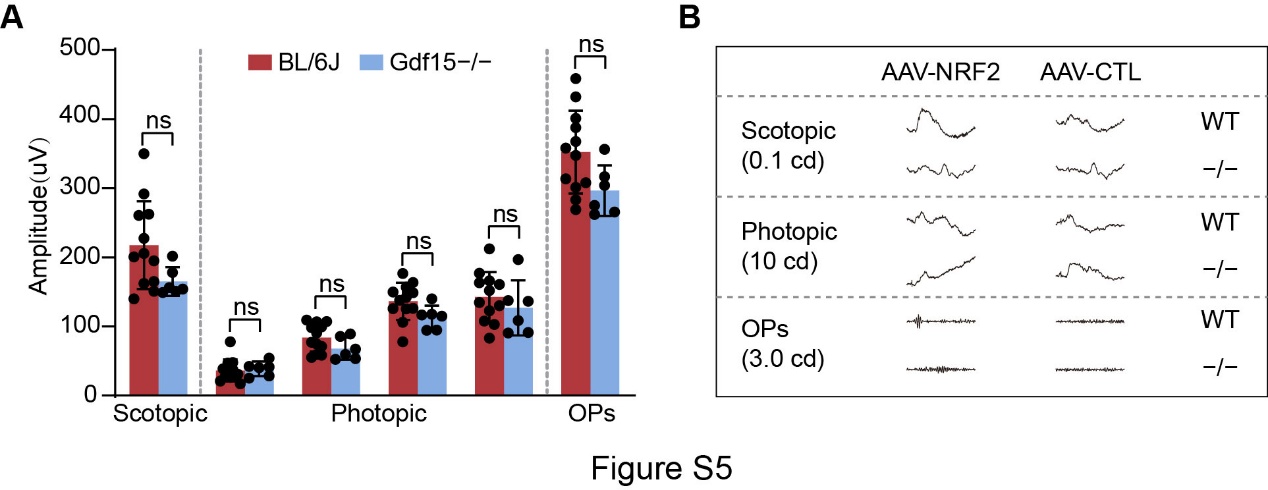

**Figure S5. ERG responses in Wild-type and Gdf15⁻/⁻ mice.**

**(A)** Quantification of baseline electroretinography (ERG) responses in wild-type (WT) and Gdf15⁻/⁻ mice (n = 12 eyes per group for WT mice, n = 6 eyes per group for Gdf15⁻/⁻ mice, mean ± SD, unpaired t-test).

**(B)** Representative ERG traces from WT and Gdf15⁻/⁻ mice receiving AAV8/BEST1-NRF2 or AAV-CTL after NaIO3 injury. Scotopic responses, photopic responses, and oscillatory potentials (OPs) are shown for the indicated groups.

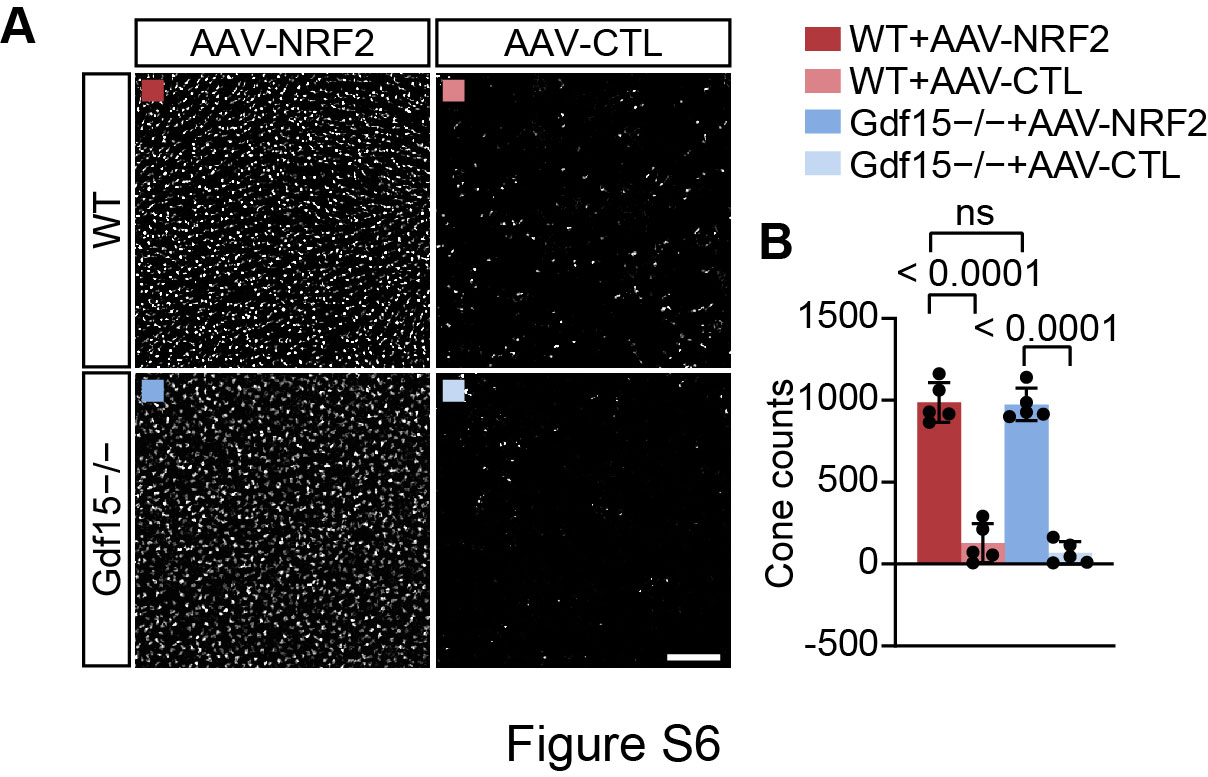

**Figure S6. Cone preservation in WT and Gdf15⁻/⁻ mice after AAV8/BEST1-NRF2 or AAV-CTL treatment.**

**(A)** Representative cone arrestin-stained retinal flat-mounts showing cone photoreceptors in WT and Gdf15⁻/⁻ mice receiving AAV8/BEST1-NRF2 or AAV-CTL following NaIO3 injury. Scale bar, 50 μm.

**(B)** Quantification of cone counts in the indicated groups (n = 5 eyes per group, mean ± SD, one-way ANOVA with Tukey’s multiple-comparisons correction).
